# RNA polymerase II initiation factors show different dynamic behaviour upon induced transcription in live cells

**DOI:** 10.64898/2025.12.19.695093

**Authors:** Attlia Oravecz, Olivier Tassy, Sascha Conic, Mustapha Oulad-Abdelghani, Nacho Molina, Laszlo Tora

**Affiliations:** Institut de Génétique et de Biologie Moléculaire et Cellulaire, 67404 Illkirch, France; Centre National de la Recherche Scientifique (CNRS), UMR7104, 67404 Illkirch, France; Institut National de la Santé et de la Recherche Médicale (INSERM), U1258, 67404 Illkirch, France; Université de Strasbourg, 67404 Illkirch, France; Present address: Deutsches Krebsforschungszentrum, Heidelberg, Germany

## Abstract

Transcription by RNA polymerase II (Pol II) requires the ordered action of general transcription factors (GTFs) forming the pre-initiation complex (PIC). How these events unfold kinetically remains unclear. Upon transcription activation, we observe coordinated recruitment of Pol II and GTFs together with chromatin decompaction. Pol II and its Ser5-phosphorylated form accumulate and persist at promoters, whereas TFIIB and TFIIF engage transiently, revealing distinct dynamic regimes. We find specific residence kinetics consistent with rapid exchange of GTFs and a more stable initiating Pol II population. A quantitative kinetic model recapitulates the temporal ordering and relative amplitudes of factor recruitment, predicts recruitment times, and links transient GTF engagement to sustained Pol II occupancy. Steady state measurements and transcription inhibition corroborate model predictions. These results uncover a dynamic hierarchy in human PIC assembly and establish a quantitative framework that connects factor exchange kinetics to the regulation of Pol II activity in living cells.

## Introduction

Most eukaryotic protein-coding genes are transcribed by RNA Polymerase II (Pol II). Initiation of Pol II transcription requires the assembly of the general transcription factors (GTFs) on core promoters to form the preinitiation complex (PIC). A plethora of mostly biochemical and structural studies have provided substantial knowledge about the structure and composition of the PIC leading to different models of its assembly ^1, 2^. According to the canonical sequential assembly model, the TATA-binding protein (TBP)-containing TFIID first recognizes, and with TFIIA, binds the core promoter. TFIIB then joins the sub-complex to support the subsequent recruitment of Pol II with TFIIF, which together form the core PIC ^3, 4, 5^. TFIIE is then recruited to the core PIC which is followed by the binding of TFIIH leading to the assembly of the holo PIC which can initiate transcription ^6, 7, 8^.

Important regulatory steps of transcription initiation and also later stages of the transcription cycle are mediated by the different phosphorylation states of the heptapeptide (Tyr-Ser-Pro-Thr-Ser-Pro-Ser) repeats in the carboxy-terminal domain (CTD) of the largest subunit of Pol II (RPB1) ^9, 10^. Current models suggest that Pol II binds to the core PIC with un-phosphorylated CTD ^11^, which has high affinity for the Mediator complex ^12, 13, 14^ important for bridging enhancers, sequence-specific transcription factors (TFs) and GTFs at the promoter ^15^. Upon the assembly of the holo PIC, the cyclin-dependent kinase 7 (CDK7) subunit of TFIIH phosphorylates Ser5 residues (pSer5) of the CTD which allows Pol II promoter escape through weakening of its interaction with mediator and enables its promoter-proximal pausing ^9, 16, 17^. Pause-release and active elongation of Pol II in turn requires CTD Ser2 phosphorylation (pSer2) by the CDK9 subunit of the positive transcription elongation factor b (P-TEFb) ^18^.

Previous studies of PIC assembly and progression of Pol II through the transcription cycle typically used structural and biochemical approaches including chromatin immunoprecipitation, footprinting and electrophoretic mobility shift assays ^5, 9, 14, 19, 20^. Though these studies provided invaluable insights into transcriptional processes and the static composition of the PIC, they were typically based on population averages and had limited temporal resolution potentially underrepresenting transient molecular interactions and masking the single cell level heterogeneity of the underlying dynamics ^21, 22^. Recent studies leveraging fluorescent microscopy and single molecule dynamics has begun to provide a better quantitative understanding of the *in vivo* kinetics of transcription initiation and its regulation.

TFIIA, TFIIB and TFIIH dynamics were analyzed *in vitro* at promoters using purified proteins ^23, 24, 25^, and more recent studies determined the single molecule dynamics of complete PIC assembly and its dependence on the Pol II CTD in living yeast cells using fluorescently tagged recombinant proteins ^26, 27^. Further single particle tracking (SPT) experiments on a single model gene using recombinantly tagged Pol II, TFIIF, TFIIE and TFIIH in yeast nuclear extracts in the presence of an activator (Gal4-VP16) suggested a branched PIC assembly pathway, alternative to the sequential assembly model, through which the PIC can first assemble into Pol II-TFIIF-TFIIE subcomplex(es) on enhancers, and TFIIH joins only after their translocation to the promoter ^28^. By photobleaching and fluorescence correlation spectroscopy assays in human cells, green fluorescent protein (GFP)-tagged TFIID and TFIIB were also shown to be highly dynamic, forming transient chromatin associations which correlated with transcription ^29^.

Single molecule dynamics of Pol II were also measured using permanently tagged fluorescent Pol II molecules at multiple un-resolved genomic locations as well as at specific single genes in mammalian cells and Pol II was shown to form transient dynamic clusters ^30, 31, 32^. In order to investigate the dynamics of endogenous proteins and their post-translational modifications in live cells, fluorescently tagged antibody-based labeling approaches were developed ^33, 34^. The power of these approaches was demonstrated by monitoring Pol II dynamics in single cells at an induced repetitive gene array ^35^ and a single HIV gene model during constant activator presence ^36^. Despite these considerable efforts, our understanding of the activator-dependent recruitment dynamics of endogenous Pol II and GTFs upon transcription activation in single human cells still remains limited.

Here, we combine fluorescent antibody-based labelling approach VANIMA ^33, 37^ with time-resolved fluorescence confocal microscopy in living cells and machine-learning-assisted image analyses to monitor the recruitment dynamics of an inducible transcription activator and endogenous TFIIB, TFIIF, Pol II and Pol II pSer5 to a gene array in live human cells. We find that Pol II (including its Ser5-phosphorylated form) accumulates and persists at activated promoters, whereas TFIIB and TFIIF display rapid, transient engagement, revealing distinct kinetic regimes within the initiation machinery. To interpret these dynamics, we develop and calibrate a quantitative kinetic model that estimates absolute molecule numbers and recruitment rates and incorporates an incoherent feed-forward inhibitory loop that explains the transient GTF behaviour; super-resolution dSTORM independently corroborates the modelled molecular abundances at steady state. Finally, pharmacological inhibition of promoter opening perturbs factor recruitment in a manner captured by the model. Together, this integrative approach provides a quantitative in vivo framework for the dynamic assembly and regulation of the human Pol II pre-initiation complex.

## Results

### Measurements of Pol II and GTF recruitment dynamics upon transcription activation

To study the recruitment dynamics of endogenous Pol II and GTFs upon transcription activation, we established a cell line based on the U2OS 2-6-3 cells ^38^, which stably express an inducible transcription activator (mCherry-tTA-ER) consisting of the tetracycline repressor (TetR) fused to a VP16 transactivation domain (tTA), a mCherry fluorescent protein and the ligand-binding domain of the oestrogen receptor (ER) (hereafter called U2OS-med cells, see Methods and Supplementary Fig. 1). Upon 4-hydroxytamoxifen (4-OHT) addition, the mCherry-tTA-ER activator enters the nucleus and binds to the tetracycline response element (TRE) containing 96 copies of specific tetO binding sites upstream of a minimal CMV promoter on a ∼200 copy gene array integrated into a single genomic position of U2OS 2-6-3 cells (Fig. 1a-b activator, Supplementary Fig. 1e).

**Fig. 1.**
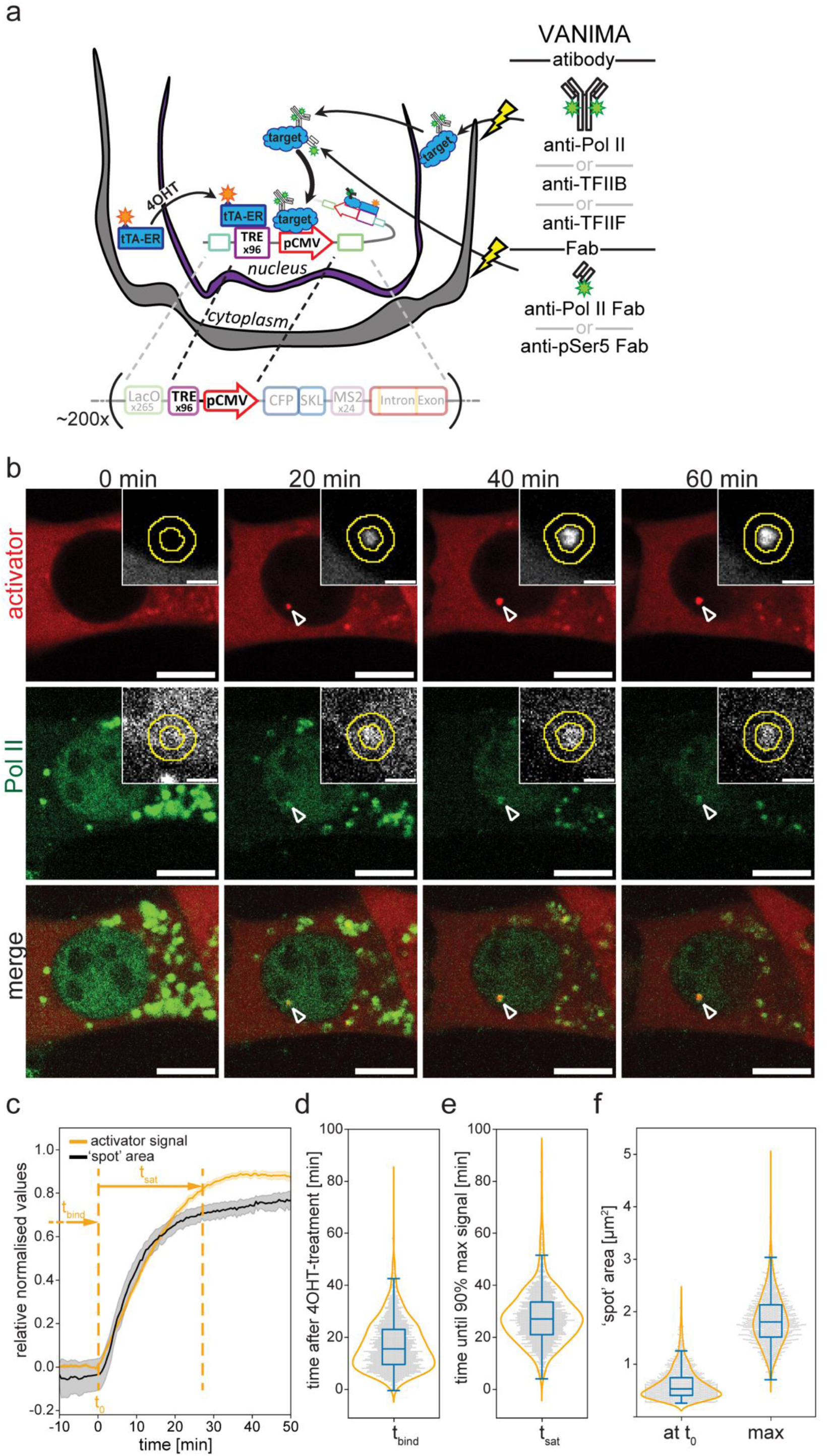
Experimental setup to measure activated recruitment dynamics of GTFs and Pol II, and characterization of the binding dynamics of the activator. **a** Schematic representation of the experimental setup. U2OS-med cells harbor a ∼200 copy repetitive tandem gene array integrated at a single locus, and express the inducible mCherry-tagged transcription activator tTA-ER in the cytoplasm. Fluorescently labelled specific antibodies targeting either Pol II, TFIIB or TFIIF, or Fabs targeting Pol II or Pol II pSer5 are used to label these endogenous factors by VANIMA ^33^. 4OHT-induced nuclear translocation of tTA-ER triggers its binding to the TRE elements present in the transcription unit (as indicated), and induce the recruitment of the GTFs and Pol II to the activated CMV promoters. The recruitment dynamics of both the induced mCherry-tagged activator and the antibody-labelled endogenous factors to the transcription sites are monitored by time-resolved fluorescent confocal microscopy. **b** Representative snapshot images at the indicated time points from of U2OS-med cells that were labelled with anti-Pol II Alexa488 antibodies overnight, activated with 4-OHT and imaged for 1 h. Arrow heads show the co-localisation (merge; bottom row) of the activator (red; top row) and Pol II (green; middle row) signals at the activated transcription sites. Scale bars are 10 µm. Insets show the tracking of the signal (inner circle) and local background (outer ring) areas highlighted by the arrowheads. Inset scale bars are 2 µm. **c** Changes of average normalized activator signal ([total signal int.-{signal area*mean background int.}]*photobleaching; re-normalized to maximum) (orange) and array size (black) over time. ±3*SEmean; n=1537. Left vertical dashed line depicts the time-point of first detected activator binding (t_0_) to which the individual traces are aligned (0 min), and the right dashed line when individual activator signals reach 90% of maximum (t_sat_). **d**, **e** Distributions of (**d**) activator binding times (Δt_bind_; median=15.5 min [95% CI 14.5-16.0]) and (**e**) saturation times (Δt_sat_; median=27 min [95% CI 26.5-27.7]). **f** Comparison of the largest area of the activator’s signal during activation (max; median=1.804 µm^2^ [95% CI 1.782-1.835]) and its area of signal at the first detected activator binding time (at t_0_; median=0.530 µm^2^ [95% CI 0.520-0.541]).

In order to measure the activator-induced recruitment dynamics of Pol II, GTFs and pSer5 Pol II CTD we performed VANIMA ^33^ to endogenously label these molecules by electroporating *U2OS-med* cells with Alexa Fluor 488 (A488)-labelled antibodies (Fig. 1a). We used previously published full length purified mouse monoclonal antibodies (mAbs) specific for TFIIB (anti-TFIIB) ^39^, the RAP74 subunit of TFIIF (anti-TFIIF) ^40^ or the CTD of the RPB1 subunit of RNA Pol II (anti-Pol II) ^41, 42, 43, 44, 45, 46^. To detect the post-translational Serine 5 phosphorylation of the hepta-peptide repeats of RPB1 CTD (pSer5), we generated a mouse monoclonal antibody specifically recognising this modification (Supplementary Fig. 2), and used the antigen-binding fragment (Fab) of this antibody (anti-Pol II pSer5 Fab), along with a control Fab of anti-Pol II (anti-Pol II Fab), both of which can freely diffuse in to the nucleus ^33^. Twenty-four hours post-electroporation we treated the cells with 4-OHT to induce the nuclear import and the binding of the activator to the transcription site. Activator binding and the recruitment of the antibody-labelled endogenous factors was monitored through the detection of nuclear mCherry and A488 foci, respectively, by confocal time resolved microscopy with 30 seconds/frame temporal resolution (Fig. 1b). 4-OHT induced the activator (mCherry) and Pol II, TFIIB, TFIIF or Pol II pSer5 (A488) signals at the transcription site in most cells (Fig. 1b, Supplementary Movie 1-5, Supplementary Fig. 3a-e). To monitor the corresponding signal intensity changes over time in 3 dimensions, we developed a machine-learning-assisted tracking and quantification pipeline (Fig 1b insets). This setup provided a platform for time-resolved quantification of activator-induced recruitment of endogenous Pol II and GTFs to the gene array.

### Characterization of the activator’s binding dynamics and the activated transcription foci

To characterise the activator’s binding dynamics, we first analysed the lag times (Δt_bind_) between 4-OHT-treatement and the first detection of significant mCherry signal (t_0_). This showed that the activator accumulated at the gene array with a median binding time of Δt_bind_=15.5 min after 4-OHT addition (Fig. 1c, d) indicating the time required for the activator’s nuclear import and binding to the TRE. We then calculated the median ‘saturation time’ (Δt_sat_) of the activator binding as time intervals between the t_0_ time points and the time points of 90% maximum mCherry signal intensities. This Δt_sat_ suggest that the activators occupy their available binding sites and reach equilibrium within 27 min after the first binding event (Fig. 1c, e, Table 1). We next compared the signal increase dynamics of the activator to the size increase of the foci of the gene array, and found that the two features grew with comparable kinetics over time (compare the progression of the orange and black curves in Fig. 1c), suggesting that the chromatin opening at the transcription site is proportional with the activators gradually occupying the gene array. Consistently, the ∼0.53 µm^2^ initial median size of the locus (Fig. 1f) increased more than 3.2-fold (Supplementary Fig. 3f) reaching a median of ∼1.80 μm^2^ maximum size (Fig. 1f). Assuming a generally spherical 3D structure, this corresponds to ∼6.3-fold expansion of the locus’ volume from ∼0.290 µm^3^ to 1.82 µm^3^. These results together indicate rapid activator recruitment and corresponding chromatin decompaction at the induced gene array upon activation.

**Table 1.** Summary table depicting the measured parameters in the study.

|  | Median time to signal saturation after activator binding [min] (95% CI) |  |  |  |  |  |
| --- | --- | --- | --- | --- | --- | --- |
|  | activator | Pol II | TFIIB | TFIIF | Pol II (Fab) | Pol II pSer5 |
| Fig. 1e, Fig. 2f | 27.0 (26.5-27.7) | 20 (19.0-21.0) | 13.5 (12.75-15.0) | 15.5 (13.5-17.0) | 22.0 (20.5-23.0) | 20.5 (20.0-22.0) |
|  | Median residence times [s] ±SEM |  |  |  |  |  |
| Fig. 3e | activator<br>174.3±34.5 | Pol II<br>46.8±5.8 | TFIIB<br>62.5±9.5 | TFIIF<br>37.1±10.0 | Pol II (Fab)<br>67.5±9.3 | Pol II pSer5<br>100.8±18.0 |
|  | Model-predicted median maximum number of bound molecules (95% CI) |  |  |  |  |  |
| Fig. 4g | activator<br>474.5 (447.8-500.2) | Pol II<br>26.9 (23.5-30.4) | TFIIB<br>176.1 (155.6-206.2) | TFIIF<br>70.1 (52.3-90.1) | Pol II (Fab)<br>43.5 (37.9-50.1) | Pol II pSer5<br>90.3 (81.7-98.7) |
|  | Steady-state recruitmet rate of Pol II or GTFs [1/min] (95% CI) |  |  |  |  |  |
| Fig. 4h |  | Pol II<br>10.72 (9.3-12.7) | TFIIB<br>8.2 (6.8-9.3) | TFIIF<br>15.29 (12.2-24.0) | Pol II (Fab)<br>10.66 (9.0-13.3) | Pol II pSer5<br>9.14 (8.3-11.0) |
|  | Steady-state recruitmet time of Pol II or GTFs [sec] (95% CI) |  |  |  |  |  |
| Fig. 4h |  | Pol II<br>5.6 (4.7-6.4) | TFIIB<br>7.3 (6.4-8.8) | TFIIF<br>3.9 (2.5-4.9) | Pol II (Fab)<br>5.6 (4.5-6.7) | Pol II pSer5<br>6.6 (5.5-7.3) |
|  | Recruitmet time of one Pol II or GTF per activator molecule [min] (95% CI) |  |  |  |  |  |
| Suppl. Fig. 5f |  | Pol II<br>84.0 (71.0-96.6) | TFIIB<br>109.9 (96.6-132.0) | TFIIF<br>58.9 (37.6-73.8) | Pol II (Fab)<br>84.6 (67.6-99.9) | Pol II pSer5<br>98.6 (82.0-109.1) |

### Pol II and GTFs show different recruitment dynamics upon transcription activation

We next compared the recruitment dynamics of Pol II, TFIIB and TFIIF together with the dynamics of pSer5 of Pol II upon induced transcription (Fig. 2). The lack of apparent shift of their average recruitment curves relative to the activator’s t_0_ time points suggests that these factors bind the transcription site short after the activator and within the temporal resolution of the assay (Fig. 2a-f). However, comparison of their first detectable binding to that of the activator’s (t_0_) revealed that Pol II and Pol II pSer5 accumulation follows activator binding with a short, but significant delay (median Δt for Pol II = 2 min, Pol II Fab = 1 min, Pol II pSer5 = 1 min) (Supplementary Fig. 3g, h). Furthermore, the binding curves also suggest that these factors reach maximum occupancy of the gene array within 20 min after activator binding. Interestingly, while in case of Pol II and Pol II pSer5 the A488 signal remains relatively stable after reaching the maximum (Fig. 2d, e), with a small decrease of the Pol II when measured with the full-length antibody (Fig. 2a), TFIIB and TFIIF signals after reaching the maximum show a marked decay (Fig. 2b-c, Supplementary Fig3. a-b, Supplementary Movie 2-3). To better characterise the different dynamics of Pol II and GTFs, we compared their median saturation times which revealed a comparable 20 and 22 min Δt_sat_ for Pol II measured by either the full-length antibody or fab, respectively, and a 20.5 min Δt_sat_ for Pol II pSer5. Interestingly, TFIIB and TFIIF showed significantly lower 13.5 and 15.5 min Δt_sat_, respectively, compared to Pol II (Fig. 2f, Table 1). These findings indicate that Pol II molecules are 20-26%, while TFIIF and TFIIB molecules are 43-50% faster than the activator to fill up the available space on the gene array. This is consistent with the activators having the most binding sites (96 per transcription unit), while only few Pol II molecules and one PIC can participate in processing one transcription unit. Altogether, these data suggest differential recruitment dynamics when comparing Pol II with TFIIB or TFIIF after transcription activation.

**Fig. 2.**
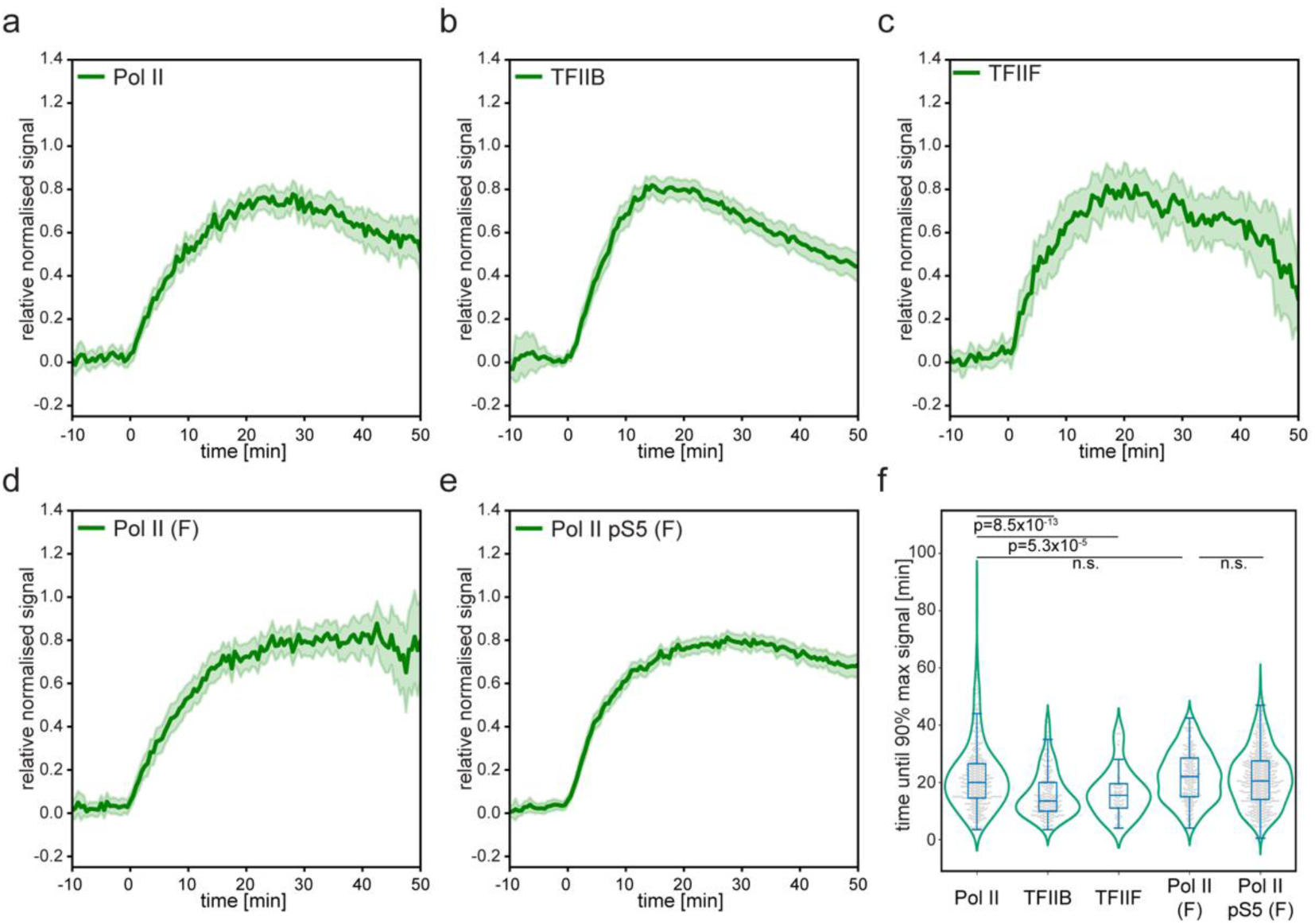
Pol II and GTFs show differential recruitment dynamics in response to transcription activation. a-e. Recruitment plots of (**a**) anti-Pol II, (**b**) anti-TFIIB, (**c**) anti-TFIIF, (**d**) anti-Pol II (Fab) and (**e**) anti-Pol II pSer5 (Fab). Changes of average normalized signals ([total signal int.-{signal area*mean background int.}]*photobleaching; re-normalized to maximum) over time aligned to t_0_= 0 min (activator binding time, see Fig 1c). For (**a**-**e**) n=412, 246, 83, 233, 563, respectively. **f** Combined violin-box-swarm plots comparing the distributions of saturation times after activator binding of the indicated factors. median [min] and (95% CI) values for Pol II: 20.0 (19.0-21.0); TFIIB: 13.5 (12.75-15.0); anti-TFIIF: 15.5 (13.5-17.0); anti-Pol II (Fab): 22.0 (20.5-23.0); anti-Pol II pSer5 (Fab): 20.5 (20.0 22.0). p-values were calculated with Holm-Šidák-correction from a post-hoc Dunn test after a significant Kruskal-Wallis test (F=101.38, p=5×10^-21^).

### Pol II and GTFs have comparable residence time during steady-state transcription

After characterizing the recruitment dynamics of the activator, Pol II, and GTFs, we next asked how long each factor remains bound at the transcription site during steady-state activation. We used fluorescence loss in photobleaching (FLIP), in which a distant nuclear region (outside the focus) is repeatedly bleached to deplete the freely diffusing fluorescent pool. Fluorescence at the transcription focus then diminishes only when bound molecules unbind and are replaced by molecules arriving from this now-bleached pool; thus, the rate of signal loss reports the typical binding duration (residence time). We established steady state by treating U2OS-med cells with 4-OHT for 24 h and monitored focus intensity while delivering brief, high-intensity laser pulses (Fig. 3a,b; Supplementary Fig. 4a–e). Under these conditions, the activator signal decayed slowly, consistent with stable binding, whereas Pol II, TFIIB, and TFIIF decayed more rapidly and to a similar extent, indicating more transient association. Pol II-pSer5 showed an intermediate decay profile, suggesting dynamics distinct from both the activator and total Pol II.

**Fig. 3.**
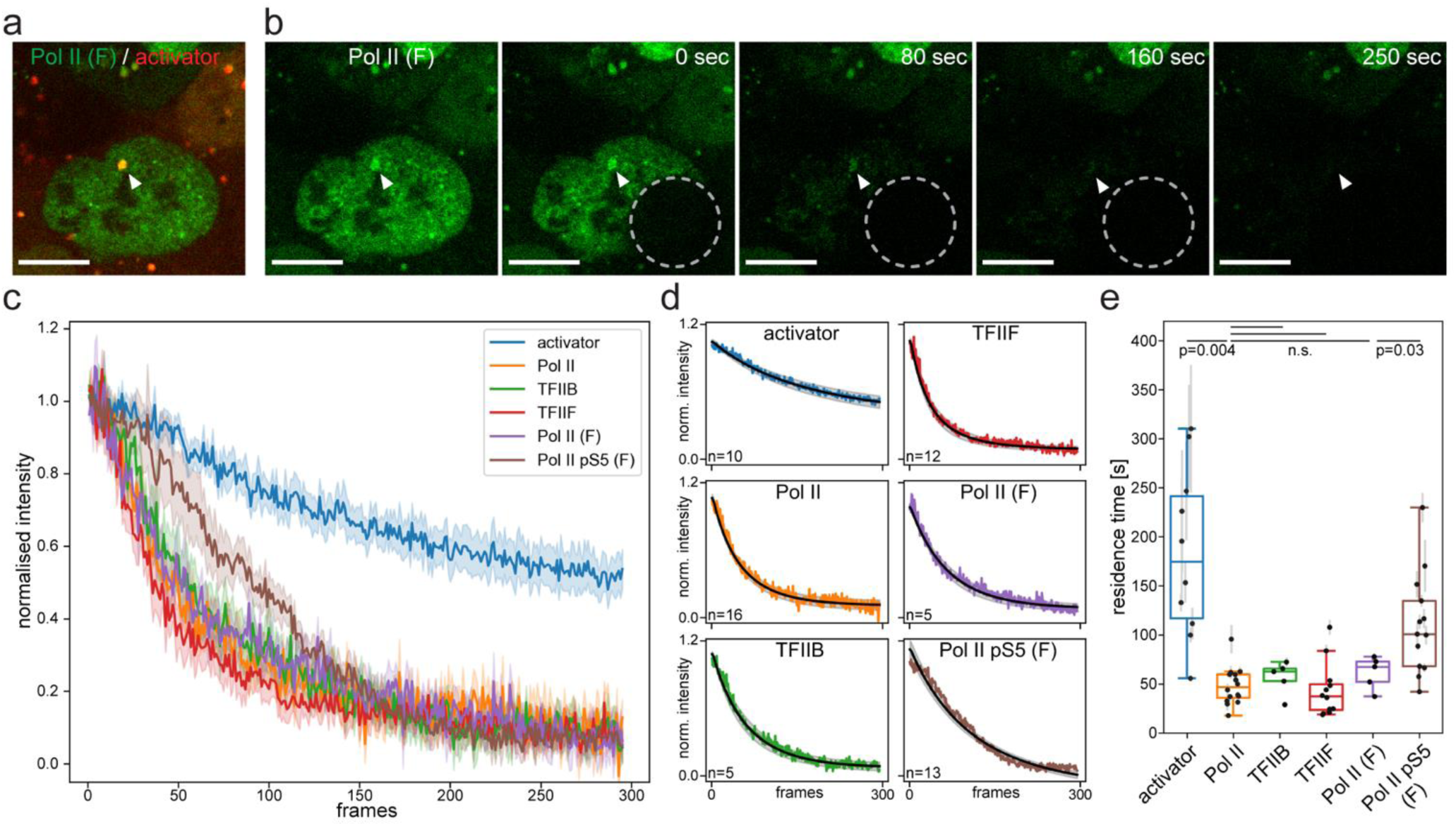
Different residence times of the activator, Pol II and GTFs during steady state transcription. a-b. Example images of the FLIP assay. U2OS-med cells were labelled with anti-Pol II Fab Alexa488 (green) and treated with 4OHT overnight. FLIP was performed with a confocal spinning disc microscope using repeated 726 ms 491 nm laser photobleaching pulses. Scale bars are 10 µm. **a** Overlay of the green and red channels. Activator signal is shown in red. Arrowhead indicate the co-localization of Pol II Fab and activator signals at the transcription site. **b** Example snapshots at select time points from the FLIP assay of the cell shown in (**a**). Left image show the first time point before photobleaching. Images taken at the indicated time points after the photobleaching pulses are also shown. Dashed circles outline the photobleached area. **c** Comparison of the average FLIP decay curves (±SEM) of the indicated factors. **d** Average FLIP decay curves from (**c**) and the corresponding average fitted single exponential functions (black curves; ±SEM) for each factor. **e** Distribution of the residence times (1/k) derived from the fitted dissociation rate constants in (**d**). median±SEmedian [sec] 1/k for activator: 174.3±34.5; anti-Pol II: 46.8±5.8; anti-TFIIB: 62.5±9.5; anti-TFIIF: 37.1±10.0; anti-Pol II (Fab): 67.5±9.3; anti-Pol II pSer5 (Fab): 100.8±18.0. Error bars: relative errors of the model’s fit for each decay curve. p-values were calculated with Holm-Šidák-correction from a post-hoc t-test after a significant one-way ANOVA (F=14.7, p=3.7×10^-9^).

To estimate residence times from these decays, we fitted each trace with a single-component model appropriate for a kinetically uniform bound population and, as a control, also fitted a two-component model to test for coexisting subpopulations with distinct lifetimes. Across all factors, the single-component model produced residence-time estimates with lower relative errors than the two-component alternative (Fig. 3d; Supplementary Fig. 4f). We therefore report residence times from the single-component fits and conclude that a single dominant binding population suffices to explain the FLIP observations. This analysis showed that the activator had the longest median residence time (174.3 ± 34.5 s), consistent with relatively stable DNA binding and in line with prior reports of stably bound transcription factors ^22^. Pol II, TFIIB, and TFIIF exhibited shorter and comparable median residence times (37.1–67.5 s), indicative of transient engagement and repeated association with the array (Fig. 3d, e; Table 1). Given the ∼3.3 kb reporter, the ∼47 s residence of Pol II corresponds to an apparent elongation rate of ∼4.2 kb min⁻¹, consistent with previous estimates. Notably, Pol II-pSer5 displayed a significantly longer median residence (100.8 ± 18.0 s) than total Pol II, consistent with its role during initiation and promoter-proximal pausing ^14, 47^.

Pol II molecule s are heterogeneously distributed in the nucleus, forming dynamic clusters that are thought to be concentrated sites of transcription often referred to as condensates, hubs or transcription factories ^48^. These endogenous Pol II clusters appear as dispersed nuclear foci on confocal images of live *U2OS-med* cells labelled with Pol II-specific fluorescent antibodies (Supplementary Fig. 4b, e). To compare the kinetic properties of Pol II within endogenous clusters and at the induced gene array, we measured signal decay from these foci using Pol II FLIP assays (Supplementary Fig. 4b). The average decay curves were similar between the activated array and endogenous Pol II clusters (Supplementary Fig. 4g). Furthermore, fitting single-exponential functions to the FLIP decays yielded median residence times that did not differ significantly between the array (46.8 ± 5.8 s) and endogenous clusters (55.1 ± 7.1 s) (Supplementary Fig. 4h,i).

Altogether our data suggest that in steady-state equilibrium of sustained transcription activation the mCherry-tTA-ER activator stably binds its target sites with a nearly 3 min residence time. The PIC components Pol II, TFIIB and TFIIF stay at the activated gene array more transiently, with a 0.5-1 min residence time, which is comparable to the residence time of Pol II molecules at endogenous clusters, suggesting that the recruitment of Pol II through the assembly of the PIC at the gene array is reflective of the physiological processes of transcription initiation.

### Mathematical model predicts Pol II and GTF absolute molecule numbers and recruitment times during transcription activation

To estimate kinetic parameters from the live-cell data, we built a two-equation model that links activator import and binding to factor recruitment at the array (Supplementary Fig. 5a). Briefly, the model tracks the number of bound activator molecules and the number of bound Pol II or a given GTF, with first-order unbinding terms for each. Activator availability in the nucleus grows with an import term (with a short lag aligned to the observed t₀), after which activator binds to a finite number of sites; Pol II/GTF recruitment is proportional to both the amount of bound activator and the remaining recruitment capacity at the array. Exit rates for Pol II and GTFs are fixed by FLIP-derived dissociation constants, so the model learns recruitment rates from the activation movies while being anchored by independent residence-time measurements (Supplementary Fig. 5a; Fig. 3d–e).

Because the imaging signals are expressed in arbitrary fluorescence units, we first implemented a calibration step to convert them into absolute molecule numbers for the activator, Pol II, and GTFs. For each trace, we smoothed the temporal trajectories and quantified the local fluctuations around the mean signal. We then assumed that these fluctuations are primarily dominated by Poisson noise, reflecting stochastic variations in the small number of molecules bound to the array, particularly at the onset of activation. Under this assumption, the variance should scale linearly with the mean intensity, allowing us to determine a calibration coefficient that converts fluorescence units into molecule counts (see Methods).

We applied the model to extract and compare the kinetic parameters of Pol II and GTF recruitment upon activator induction. The model captured well the dynamics of activator binding and the accumulation of Pol II and Pol II pSer5 molecules, as shown by close fits to the calibrated recruitment curves (Supplementary Fig. 5b) and low fitting errors. However, it failed to reproduce the TFIIB and TFIIF data (Supplementary Fig. 5b, c) that resembles a temporal pulse response (Fig 2b, c, Supplementary Fig. 5b, c). To address this, we incorporated an activator-induced type-1 incoherent feed-forward loop (I1-FFL) into the original model ^49, 50^, in which the activator directly promotes GTF and Pol II recruitment while also triggering an inhibitory mechanism (I) acting on the same factors (Fig. 4a). The upgraded model retained good fits to Pol II and Pol II pSer5 and showed greatly improved fits to both the TFIIB and TFIIF data (Supplementary Fig. 5c, Fig. 4b-f).

**Fig. 4.**
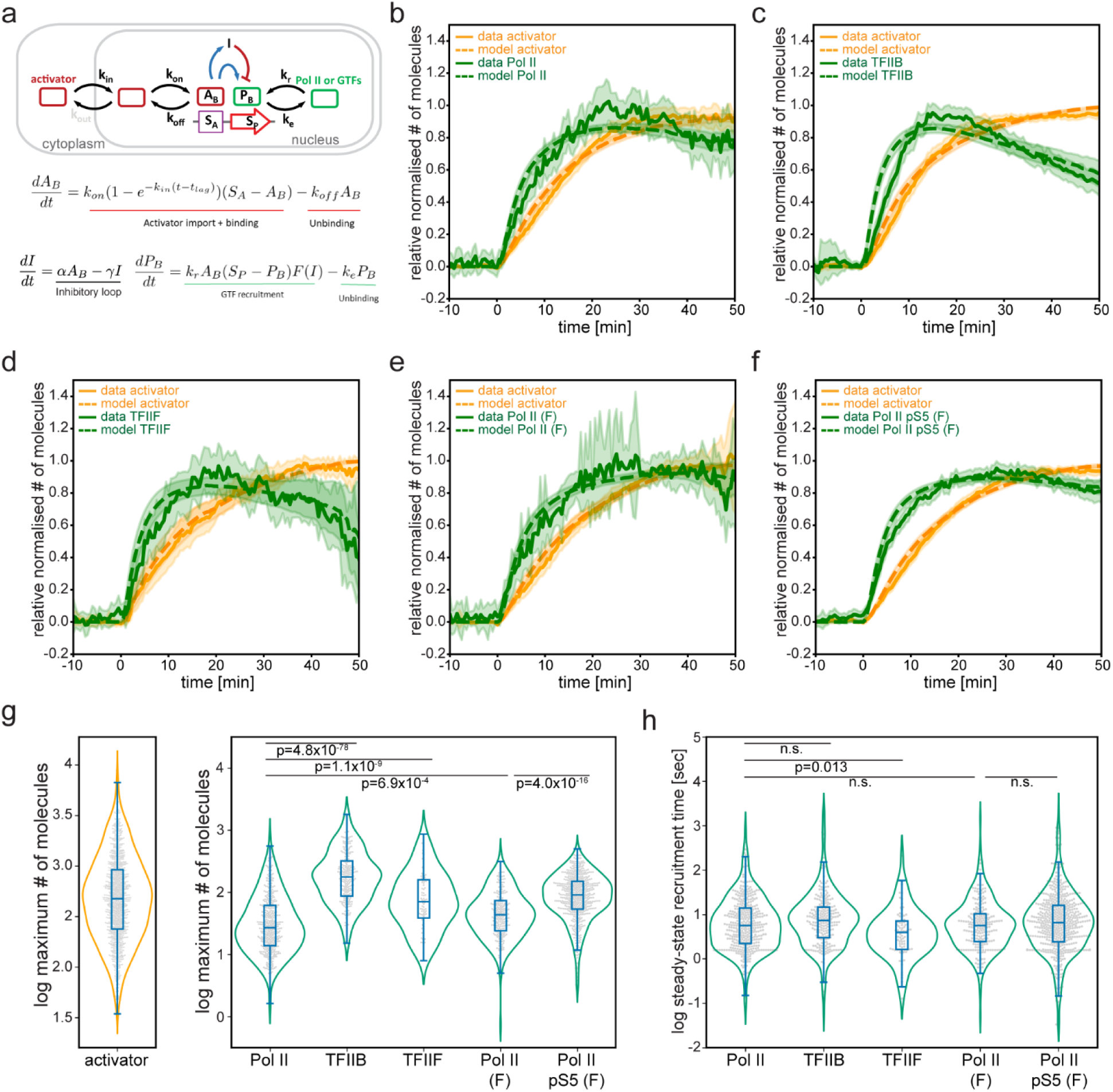
Two-state kinetic model estimates the number of molecules and recruitment times of activator, Pol II and GTFs. **a** Updated recruitment model for activator, Pol II and GTFs including an incoherent feedforward inhibitory loop. **b-f.** Calibrated binding curves and model fits for Pol II (**b**), TFIIB (**c**), TFIIF (**d**), Pol II Fab (**e**), Pol II pSer5 Fab (**f**) (green) and activator (orange) using the updated recruitment model. Solid and dashed lines show average normalized calibrated data (±3xSEM) and the average fitted models (±3xSEM), respectively. **g** Distributions of model-predicted maximum numbers of bound molecules of activator (left) and the indicated factors (right). median and (95% CI) values for activator: 474.5 (447.8-500.2); Pol II: 26.9 (23.5-30.4); TFIIB: 176.14 (155.6-206.2); TFIIF: 70.1 (52.3-90.1); Pol II (Fab): 43.5 (37.9-50.1); Pol II pSer5 (Fab): 90.3 (81.7-98.7). p-values were calculated with Holm-Šidák-correction from a post-hoc Dunn test after a significant Kruskal-Wallis test (F=431.69, p=3.95×10^-92^). **h** Distributions of model-predicted steady-state recruitment times of the indicated factors. median [sec] and (95% CI) values for Pol II: 5.6 (4.7-6.4); TFIIB:7.3 (6.4- 8.8); TFIIF: 3.9 (2.5-4.9); Pol II Fab: 5.6 (4.5-6.7); Pol II pSer5 Fab: 6.6 (5.5-7.3). p-values were calculated with Holm-Šidák-correction from a post-hoc Dunn test after a significant Kruskal-Wallis test (F=18.11, p=0.001).

We then used the refined model to estimate number of molecules recruited to the gene array during activation. To compare the activator, Pol II and the GTFs, we analysed the distributions of the maximum numbers of molecules bound to the gene array predicted from the calibrated fits (Fig 4b-g). We found that the median of the maximum numbers of activator molecules binding the gene array was 474.5±27.44, with some transcription sites recruiting more than 3000 activator molecules, while some others only less than 100 (Fig. 4g, left). Given the nearly 20000 potential activator binding sites on the gene array (96×200 TetO sites) (Fig. 1a), this result suggests that only a fraction of these sites is occupied by activator molecules. The model predicted a 26.9±5.13 median maximum total Pol II molecule number on the activated gene array when measured with the full-length antibody, and interestingly this number was 6.5-times higher for TFIIB (176.14±23.56) and 2.6-times higher for TFIIF (70.1±21.03) (Fig. 4g, right). Furthermore, when comparing data measured using fabs, we observed a 2-fold higher accumulation of Pol II pSer5 (90.3±4.44) molecules compared to total Pol II (43.5±4.77). Notably, according to these model predictions, fabs detect 1.6-times more Pol II molecules than the full-length antibody.

Next, we investigated the recruitment dynamics of Pol II and GTFs upon activator binding by comparing their model-derived recruitment rates (k_r_) and the corresponding recruitment times (1/k_r_). It is important to note, that our recruitment time is defined per bound activator molecule; in other words, how long it takes for one Pol II or GTF molecule to be recruited to the gene after one activator molecule binds its target site. For Pol II, the model predicted recruitment times of ∼84 minutes (k_r_=0.0119±0.0041 and 0.0118±0.0075 min^-1^ for the full-length antibody and the fab, respectively) and ∼99 min for Pol II pSer5 (k_r_=0.0101±0.0086 min^-1^) (Supplementary Fig. 5d, Table 1). Interestingly, while the model predicted a TFIIB recruitment time not significantly different from Pol II (though slightly longer at ∼110 min with kr=0.0091±0.0064 min^-1^), it revealed a significantly shorter recruitment time for TFIIF of ∼59 min (kr=0.017±0.0099 min^-1^) (Supplementary Fig. 5d; Table 1).

Although the per-activator recruitment times derived from the model appear long (e.g., ∼84 min per Pol II molecule for a single bound activator), this parameter reflects the waiting time per activator molecule. In vivo, many activators bind in parallel and their effects add up. As soon as the first activator binds (t₀), the number of bound activators rises and reaches saturation by ∼27 min; within our 30 s resolution, Pol II and GTF signals increase in parallel and saturate within ∼13–22 min (Fig. 1c, e; Fig. 2f; Fig. 4b–f). These observations indicate that aggregate recruitment is much faster than the per-activator baseline. Concretely, if hundreds of activators are bound at a time, the effective recruitment interval scales down from minutes to seconds (e.g., ∼84 min per activator translates to ∼seconds per event when ∼10²–10³ activators are engaged), which is exactly the regime captured by the fits to the accumulation curves.

To formalize this multiplicity effect, we rescaled the per-activator rate by the median steady-state number of bound activators (≈ 900; Supplementary Fig. 5e) and obtained steady-state recruitment times on the order of seconds: ∼5.6 s for Pol II, ∼6.6 s for Pol II-pSer5, ∼7.3 s for TFIIB (not significantly different from Pol II), and ∼3.9 s for TFIIF (significantly faster) (Fig. 4h; Table 1). Thus, while a single transcription factor would be an inefficient recruiter on its own, the collective action of many bound activators yields rapid, physiologically plausible recruitment, helping explain why genes typically employ multiple regulatory elements and TFs to achieve timely activation.

### Super resolution microscopy estimates the total number of molecules binding the transcription site and confirms model predictions

The gene array contains nearly 20000 potential biding sites for the activator and harbours approximately 200 target gene copies (Fig. 1a). Unexpectedly, our dynamic model predicted the binding of only a 400-500 of activator and 30-40 Pol II molecules to the activated gene array on average (Fig. 4g). Therefore, we wanted to verify whether our model would correctly estimate the binding of of Pol II. Thus, we sought to measure the number of activator and Pol II molecules recruited to the gene array using super resolution single molecule microscopy. We stained one day 4-OHT-treated U2OS-med cells with Alexa647-labelled antibodies specific for the activator or Pol II and performed direct stochastic optical reconstruction microscopy (dSTORM). We then applied an analyses pipeline of segmentation, thresholding and calibration on the processed super resolution images to quantify single molecules. The mCherry signal of the activator foci were used to delineate the transcription site at which the activator and Pol II single molecules were quantified compared to random nuclear regions of same sizes. These analyses revealed the presence of 598±148 activator molecules per µm^2^ at the transcription site (Fig. 5a), corresponding to 1688±408 total activator molecules on average at the gene array (Supplementary Fig. 6a), both significantly higher than at the average activator presence at the control regions. We measured 52±15 Pol II molecules per µm^2^ at the transcription site (Fig. 5b), corresponding to 177±29 total Pol II molecules on average at the gene array (Supplementary Fig. 6b).

**Fig. 5.**
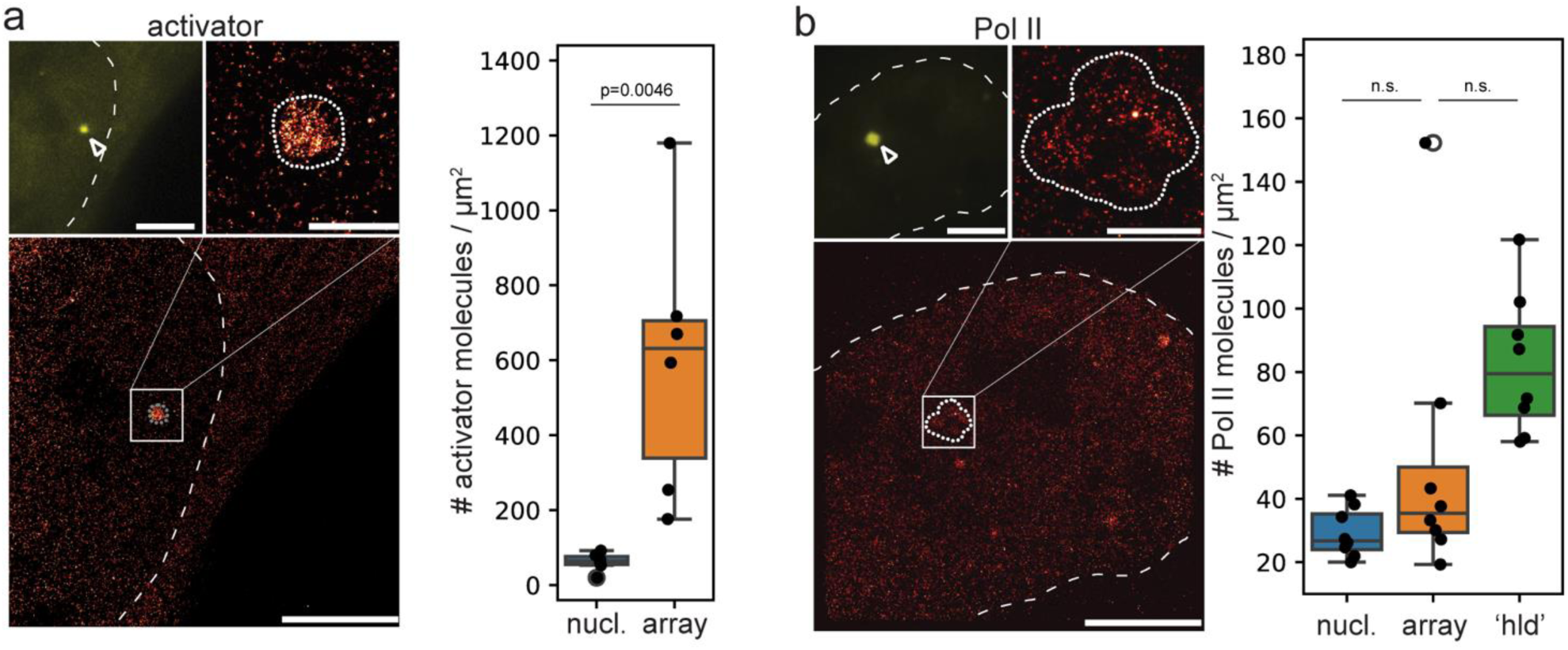
Quantification of activator and Pol II molecules bound to the active transcription site confirm model predictions. a-d. Super resolution imaging of activator (a) and Pol II (b) immunostained with specific Alexa647-labeled antibodies in U2OS-med cells treated with 4OHT overnight. Images on the left are examples of dSTORM images. Top left image: activator mCherry fluorescence showing the location of the gene array (arrowheads). Scale bar 10 µm. Bottom image: Super resolution image of Alexa647 from the same field of view as the top left image. Scale bar 10 µm. Top right image: enlarged view of the indicated regions from the bottom images. Scale bar 1 µm. Dashed lines mark the nuclei, dotted lines delineate the estimated positions of the transcription sites. Box plots on the rights show quantifications of the of Alexa647 localizations normalized by the size of the transcription site at the indicated nuclear regions. For (a), measurements at the gene array (array) and at random nuclear regions of the same size as the gene array (nucl.) are shown (n=6). mean±SEM activator array: 598.41±147.6; activator nucl.: 62.01±10.16. *p*-value is calculated from a Student’s *t*-test For (b), measurements at high local density nuclear regions (hld) are also shown (n=8) mean±SEM Pol II array: 51.65±15.34; Pol II hld.: 82.54±7.89; Pol II nucl.: 29.2±2.74. *p*-values were calculated from Student’s *t*-tests with Bonferroni correction.

The comparison with the model-predicted molecule numbers (Fig. 4g, Table 1) revealed that the super resolution image quantifications detected 3.55-times more activator (475 vs. 1688) and 6.57-times more Pol II (27 vs. 177) molecules at the activated gene array. Note however, that the model predictions were based on dynamic data obtained within one hour after transcription induction, while the super resolution microscopy was performed when the transcription from the gene array was induced to steady-state equilibrium, which may at least partly explain the differences between the model predicted and the measured molecule numbers. Nevertheless, both the model predicted and the measured activator and Pol II molecule numbers were less than the number of available binding sites and the number of genes on the gene array, suggesting that only a portion of the nearly 20000 tetO sites are bound by activators, and only a subset of the approximately 200 genes of the gene array are activated at the same time.

To investigate whether Pol II recruitment at the gene array is comparable to Pol II molecule numbers at endogenous Pol II clusters/foci, we identified ‘high local density’ (hld) nuclear regions as features characteristic of endogenous clusters and quantified the number of Pol II molecules in these regions. We found 83±8 Pol II molecules per µm^2^ at the predicted endogenous clusters, corresponding to an average of 337±52 total Pol II molecules (Fig. 5b, Supplementary Fig. 6b) per cluster. This shows a two-fold difference in the number of Pol II molecules at the 1.6-times more dense endogenous clusters compared to the gene array (Supplementary Fig. 6b).

### Transcription inhibition perturbs the model-predicted recruitment dynamics of Pol II and TFIIB

To further investigate if and how the dynamics of Pol II, Pol II pSer5 and TFIIB is altered during perturbed transcription, and to evaluate if our mathematical model can capture these changes we pre-treated antibody-labelled *U2OS-med* cells with Triptolide (TPL; 5 µM) for 2 h before transcription activation and time laps imaging. TPL inhibits transcription initiation by interfering with promoter DNA melting through inhibiting the ATPase activity of the XPB helicase subunit of TFIIH ^51^. We first evaluated if TPL treatment affected the binding of the activator by comparing the combined average activator binding curves of mock-and the TPL-treated samples from the three sets of anti-Pol II,-Pol II pSer5 and –TFIIB labelled samples, which revealed very similar activator binding dynamics between the two conditions (Fig. 6a, left). Furthermore, comparisons of the activator binding times and the maximum activator signals revealed mostly statistically insignificant differences between the control and the TPL-treated samples (Fig. 6a, right and Supplementary Fig. 7a-b). These results suggest that TPL does not have major effect on activator binding time and dynamics, nor it affects significantly the number of activator molecules binding to the transcription site enabling the study of TPL’s impact on transcription downstream of activator-binding.

**Fig. 6.**
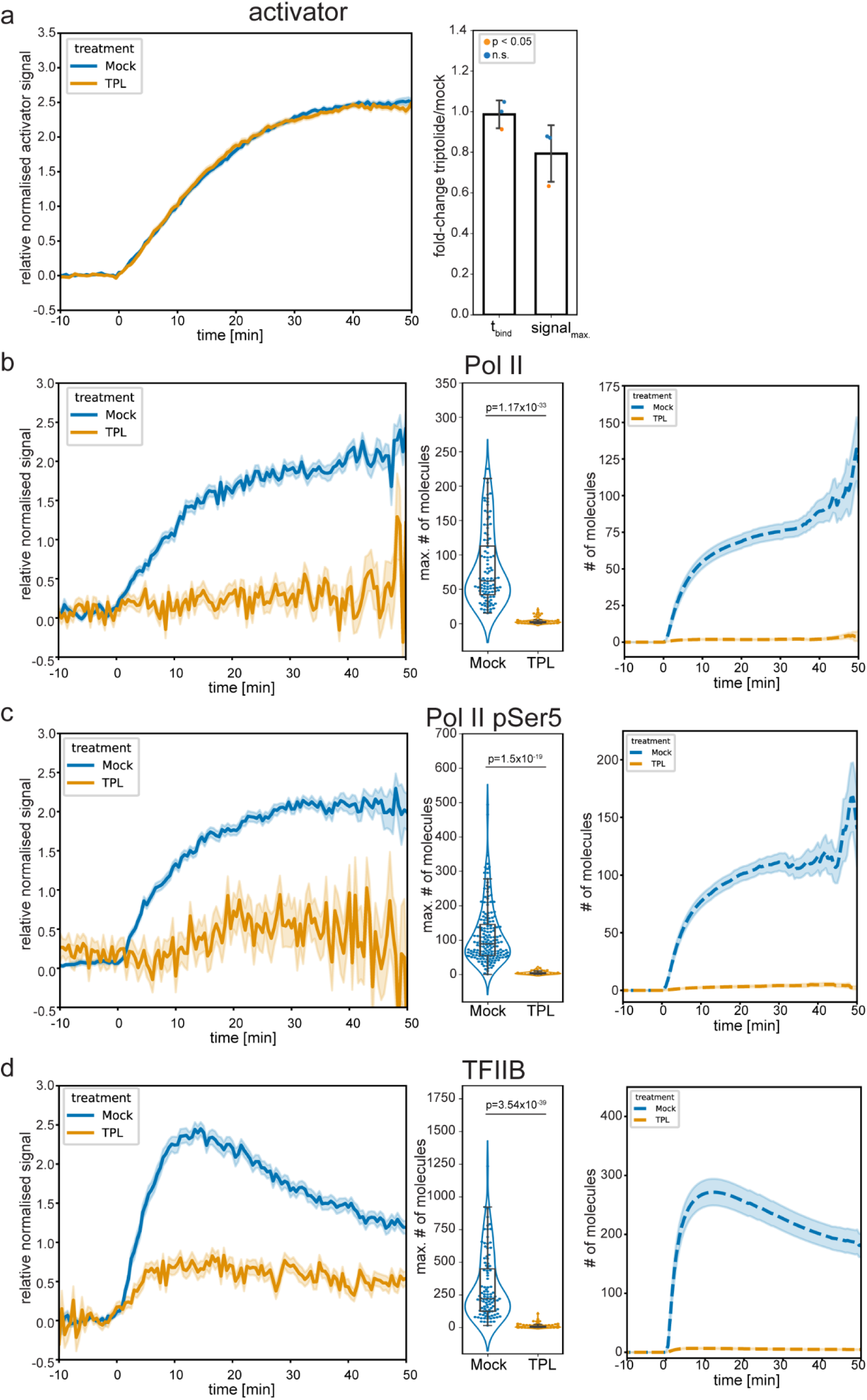
The effect of transcription inhibition on the dynamics of Pol II, Pol II pSer5 and TFIIB. Endogenous Pol II, Pol II pSer5 or TFIIB of U2OS-med cells were labeled with VANIMA, and the cells were treated with mock (DMSO) or 5µM triptolide (TPL) for two hours prior transcription activation and imaging. **a** TPL does not have a major effect on activator binding. Left: relative average normalized activator recruitment curves ([total signal int.-{signal area*mean background int.}]*photobleaching; re-normalized to SD) (±SEM). Right: Average fold-change (FC) of activator binding times (t_bind_) and maximum activator signals (signal_max_) were calculated by comparing the medians of single distributions from three different sets of antibody-labelled samples. Δt_bind_=0.99±0.07; Δsignal_max_=0.79±0.14. Error bars: SD. Single data points are color-coded according to the statistical significance of each of the three comparisons. (see Supplementary Fig. 7a, b for details). **b**-**d** The effect of TPL on Pol II (**d**), Pol II pSer5 (**e**) and TFIIB (**d**) recruitment after transcription activation. Left panels show relative average normalized recruitment curves (re-normalized to SD) (±SEM), middle panels show the distributions of model-predicted maximum number of bound molecules and right panels show fitted models (without re-normalization) (±SEM).

We therefore compared the activator-induced accumulation of Pol II, Pol II pSer5 and TFIIB at the transcription site in the presence of TPL. The average binding curve of Pol II showed a complete block of Pol II recruitment to the gene array (Fig. 6b, left), in agreement with previous results reporting the strong and fast inhibitory effect of TPL on steady-state transcription ^36^. This strong effect might also be explained by decreased Pol II protein stability induced by TPL ^52^ (Supplementary Fig. 7c). In accordance, our model predicted a marked decrease in the number of Pol II molecules binding the transcription site (median 2.07±0.47 vs 61.86±6.66) and also showed a flat recruitment profile (Fig. 6b middle and right). Similarly, to Pol II recruitment, the average binding curve of Pol II pSer5 indicated that TPL strongly decreased the appearance of CTD Ser5 phosphorylation at the gene array upon transcription activation (Fig. 6c, left), together with a marked decrease of the model-predicted number of Pol II pSer5 molecules (median 3.98±1.01 vs 88.64±7.88) and a non-increasing model profile (Fig. 6c, middle and right). This decrease of Pol II pSer5 is likely a consequence of TPL-induced Pol II destabilization rather than a direct effect on Pol II phosphorylation. Interestingly, TPL also inhibited the recruitment of TFIIB to the transcription site as indicated by the strongly decreased TFIIB binding curves compared to mock controls (Fig. 6d, left). Accordingly, our model predicted a marked diminution in the numbers of TFIIB molecules binding to the gene array (median 4.64±1.46 vs 221.12±30.41), and a flat TFIIB recruitment profile (Fig. 6d, middle and right). Altogether, these results indicate that TPL blocks Pol II, and strongly inhibits TFIIB recruitment even when activators efficiently bind to the transcription site and that our mathematical model effectively recapitulates these changes.

## Discussion

In this study, we combined VANIMA-based endogenous labelling ^33^, time-resolved live-cell imaging, FLIP, and quantitative modelling to dissect the dynamics of TFIIB, TFIIF, Pol II and, and Pol II Ser5 phosphorylation during transcriptional activation at a multigene array. This integrative approach revealed distinct kinetic regimes among initiation factors and linked those regimes to a minimal regulatory architecture that explains the observed temporal profiles, providing insights into the non-equilibrium dynamics of transcription activation.

Transcription factors bind DNA transiently and while their reported residence or survival times are typically on the timescale of seconds, some TFs and even sub-populations of certain TFs can remain bound to their target DNA for minutes ^22, 53, 54^. The relatively long residence time (∼174 s, Fig. 3e) we measured for tTA-ER in our system places this model TF among longer-binding TFs suggesting that it acts as a more stably binding transcription activator ^22^. Interestingly, TetR alone has been suggested to reside on chromatin only on the order of seconds (non-specific binding for 5 s) ^55, 56^ indicating that the stable interaction of tTA-ER with the transcription site is likely conferred, at least in part, by the VP16 activation domain potentially through stabilizing interactions with chromatin re-modellers, transcription co-activators and the transcription machinery ^57^. Note however, that the high number of potential binding sites (about 20 000) on the gene-array may also contribute to the measured high residence time through synergistic or cooperative binding of tTA-ER, potentially resulting in higher-order structural folding.

We measured a 46.8 s residence time for endogenous total Pol II on the gene array in steady state transcription (Fig. 3e). Importantly, we found no significant difference between the Pol II residence times on the gene array and at endogenous transcription clusters (55.1 s) (Supplementary Fig. 4g-i) suggesting that the transcription machinery may operate with comparable dynamics at the gene array and endogenous transcription clusters. Since we found that a single exponential decay function fits best our FLIP data, and since the total Pol II antibody detects all Pol II forms, these Pol II residence times likely represent a bulk average of promoter-bound, initiating and elongating Pol II molecules. In support of this, the residence time of exogenous expressed Pol II fused to YFP has been estimated to be ∼54 s for initiating, and ∼32 s for actively elongating fractions by FRAP ^58^. It should be noted, that the absence of a fast-decaying Pol II fraction in our FLIP data corresponding to transiently binding Pol II with short (<10 s) residence time is likely due to our local background subtraction method during FLIP image analyses. Nevertheless, our FLIP measurements directly monitoring the endogenous initiating Pol II pSer5 molecules revealed a 100.8 s residence time, 50% higher than the corresponding total Pol II control (Fig. 3e). The longer residence time of Pol II pSer5 reflects its longer interaction with the transcription site due to transcriptional pausing. Interestingly, though a single exponential decay function fits the Pol II pSer5 FLIP data with acceptable errors (Fig. 3d, Supplementary Fig. 4f), the decay profile of the signal is distinct from the total Pol II and the other GTFs signals, and resembles more a sigmoid decay curve (Fig. 3c). This, together with the fact that a double exponential decay function did not fit the decay curve, suggest a more complex dynamic behaviour of Pol II pSer5. The presence of an unbound pool of Pol II pSer5 near the transcription site potentially originating from abortive initiations have been proposed ^36, 59^. These molecules would then be able to quickly re-integrate into the promoter PICs, and re-initiate transcription skipping the PIC assembly step. As Pol II Ser5 phosphorylation can only occur during transcription initiation, these molecules would not be replaced by the dark Pol II molecules from the nucleoplasm, and could distort the exponential decay profile by ‘buffering’ the fluorescence loss.

We measured comparable, and not significantly different residence times for total Pol II, TFIIB and TFIIF during an activator stimulated PIC assembly (46.8, 62.5 and 37.1, respectively). These findings contrasts previous single molecule measurements in yeast ^26^ reporting 2-10 s residence times for PIC components and 20-22 s for Pol II and *in vitro* single molecule TIRF assays ^25^ reporting 1.5 s residence time for TFIIB.

Our kinetic modelling approach builds on—and bridges—two traditions in live-cell transcription analysis. FRAP/strip-FRAP studies at engineered or endogenous loci fit reaction–diffusion or multi-step initiation models directly to fluorescence time courses to estimate kinetic rates for Pol II engagement and promoter escape ^58, 59^, while single-gene MS2/PP7 and single-molecule tracking studies infer promoter-state switching and factor dwell times from nascent-RNA or factor trajectories ^26, 36, 60,61^. We extend this repertoire by coupling FLIP-anchored exit rates to an ODE model that explicitly links activator nuclear entry and binding capacity to Pol II/GTF recruitment, and by introducing an incoherent feed-forward motif that parsimoniously explains the transient TFIIB/TFIIF pulses. A second departure is our calibration to absolute molecule numbers from the live movies themselves. Conceptually, this is related to FCS, which uses intensity autocorrelation to extract particle number and diffusion; here, we exploit variance–mean scaling of local fluorescence fluctuations (most informative near activation onset when copy numbers are small) to derive a conversion factor from arbitrary units to molecules. Unlike classical FCS, which requires a defined observation volume and assumes stationarity of freely diffusing species, our calibration operates at a fixed genomic focus during activation, complements FRAP/FLIP-style fits, and yields absolute occupancies that we independently cross-validate by dSTORM.

Importantly, our mathematical model that fits best the activated transcription dynamics data incorporates an incoherent feed forward loop ^49, 50^ to account for the transient binding characteristics of TFIIB and TFFIIF. This loop predicts an inhibitory mechanism acting on the binding dynamics of these two GTFs after their initial activator-induced recruitment. The molecular nature of this inhibitory pathway remains however elusive. It is possible that the robust activation of many transcription units in the gene array induces the recruitment of a large number of regulatory proteins during the first phase of the activation causing crowding of these molecules at the transcription site. This would possibly create a steric barrier for the binding of TFIIB and TFFIIF eventually leading to their decreased presence approaching equilibrium after the summit of activation. A more direct regulatory pathway could involve a kinetic proofreading mechanism elicited by a yet unknown factor downstream of the activator and PIC assembly regulating their dynamics ^62, 63^. Alternatively, it is also conceivable that TFIIB and TFFIIF recruitment is less/not needed for re-initiation during new rounds of transcription cycles by Pol II molecules under continuous activation of the loci. Nevertheless, further precise measurements will be needed to uncover the in vivo dynamic hierarchy of all PIC components, to be able to establish a live cell kinetic framework of GTFs during the regulation of Pol II activity.

## Methods

### Cell culture and inhibitor treatments

U2OS 2-6-3 cells ^38^ were kindly provided by Susan M. Janicki. These cells harbor a single integration of ∼200 copies of a gene cassette at a euchromatic site. The gene cassette is composed of 256 *lacO* repeats followed by 96 Tet operators, a CMV minimal promoter with a downstream transcription unit of a cyan fluorescent protein fused to a peroxisome targeting signal tripeptide sequence, 24 MS2 repeats, the last intron-exon module and the terminator of the human beta-globin gene ^38^. U2OS 2-6-3 cells were cultured in a humidified incubator at 5 % CO_2_ and 37°C in Dulbecco’s modified Eagle’s medium (DMEM) (4.5 g/liter glucose) supplemented with 2 mM GLUTAMAX-I, 10% tetracycline-free fetal calf serum (FCS), penicillin (100 UI/ml), and streptomycin (100 μg/ml). Triptolide (TPL, #T3652; Sigma-Aldrich) and THZ1^64^ (#HY-80013; CliniSciences) transcription inhibitor stock solution was prepared in DMSO (Sigma-Aldrich) and stored at-20°C. For live imaging TPL was first pre-diluted in 100 µl medium and then added to the cells for 2 hours before imaging at 5µM final concentration. For immunofluorescence assays cells were treated with 15 µM THZ1 for 2 h before fixations.

### Generation of *U2OS-med* cell line

To generate U2OS 2-6-3 cells stably expressing the mCherry-labelled and 4-OHT-inducible mCherry-TetR-VP16-ER activator, first the DNA sequence encoding the TetR-VP16-ER fusion was inserted in the *Kpn*I-*Bam*HI sites of the pmCherry-C3 mammalian expression vector (pmCherry-C3-tTA-ER; sequence available upon request). 6 x 1.2×10^6^ U2OS 2-6-3 cells were electroporated with 6 x 6.85 µg linearized (*AseI* digestion) pmCherry-C3-tTA-ER vector using the Neon Transfection system with the 100 µl Neon electrode tips (MPK5000 and MPK10096, Thermo Fisher Scientific). mCherry-positve cells were sorted the next day using FACSAria II SORP (BD Biosciences) with 85 µm nozzle (Supplementary Fig 1a). mCherry fluorescence was collected using the 561 nm laser line and the 600LP-610/20BP filter cube, and autofluorescence was measured using the 700LP-780/60BP filter cube. Sorted cell were allowed to recover in culture for 2 days then expanded in the presence of 700 µg/ml G418 for 15 days. Cells were harvested, and cell expressing medium levels of mCherry were enriched by three rounds of repeated re-sorting and expansion in culture for 7 days under G418 selection (Supplementary Fig 1b). The resulting cell line was named U2OS-med, and validated for medium mCherry expression by flow cytometry using LSRFortessa (BD Biosciences) (Supplementary Fig 1c). U2OS-med cells were then expanded for 36 days over 10 passages and stable mCherry expression was validated by flow cytometry using LSRFortessa at days 2, 22 and 36 (Supplementary Fig 1d).

### Antibody production

The anti-Pol II CTD pSer5 antibody was generated as previously described ^65^. Briefly, eight-week-old Balb/c female mice were injected intraperitoneally with 100 μg of ovalbumin-coupled CTD_3_ peptides, on which the 5^th^ Serine residues were phosphorylated, and 200 μg of poly(I/C) as adjuvant. Three injections were performed at 2 weeks intervals and four days prior to hybridoma fusion, mice with positively reacting sera were re-injected. Spleen cells were fused with Sp2/0.Agl4 myeloma cells as described ^66^. Culture supernatants of growing hybridoma clones were screened by ELISA using non-phosphorylated and phosphorylated CTD peptides. Specific cultures were cloned twice on soft agar and a clone specific for pSer5 CTD (19PB-2E6) was selected. The antibody was produced as culture supernatant. All animal experimental procedures were performed according to the French and European authority guidelines.

### Antibody purification

The following mouse monoclonal antibodies were generated in house: anti-Pol II is specific for the C-terminal repeat domain (CTD) of the largest subunit of Pol II (1PB-7G5; #IG-PB-7G5 Euromedex)^41, 42, 43, 44, 45, 46^; the anti-pSer5-Pol II is specific for the Pol II CTD (19PB-2E6; see above); the anti-TFIIB (2G8) for human TFIIB ^39^; anti-TFIIF is specific for the RAD74 subunit of TFIIF (2A3) ^40^; anti-ER was raised against the peptide corresponding to amino acids 578-595 of the human estrogen receptor alpha (#04-1564; Sigma-Aldrich) ^67, 68^. Antibodies were purified as previously described ^33^. Briefly, 1 ml of antibody-containing ascites was incubated with 1.2 ml of settled bead volume of pre-equilibrated Protein G Sepharose Fast Flow beads (GE Healthcare) for 2 hours at 4°C with gentle agitation. Beads were then washed on a Poly-Prep Chromatography column (Bio-R ad) with 20 column volumes of PBS and the antibody was eluted in 1 ml fractions using 0.1 M glycine (pH 2.7) and immediately neutralized with 70 to 90 μl of 1 M tris-HCl (pH 8.0). Aliquots (6.5 μl) from each fraction were analyzed by SDS–polyacrylamide gel electrophoresis (SDS-PAGE) and coomassie blue staining, and the fractions containing most of the antibodies were pooled and dialyzed in DiaEasy Dialyzer 6 to 8 kDa MWCO dialysis tube (K1013-100, BioVision) against 2 liters of PBS overnight and for 2 hours with 2 liters of fresh PBS. The antibody solution was then concentrated on Amicon Ultra-4 centrifuge filters with 50 kDa MWCO (Millipore) to 1 to 4 mg/ml in PBS.

### Preparation of antigen binding fragments (Fabs)

Fabs of anti-Pol II and anti-pSer5-Pol II were prepared using 100 µl settled bead volume of immobilized papain agarose beads (#20341; Thermo Scientific) pre-washed three times with PBS, then incubated with 400 µl 1 µg/µl antibody solution in PBS containing 1 mM TCEP (HR2-801; Hampton Research) for 3 h in a heating bock at 37°C shaking at 800 rpm. The SN was collected using 800 µl centrifuge columns (#89868; Pierce), and the beads were washed once with 400 µl PBS, and the flow-through was combined with the SN. Undigested antibodies and Fc fragments were removed by adding the SN to ∼400 µl settled bead volume of ProteinA Sepharose beads (P9424-5ML; Sigma-Aldrich) pre-washed with PBS three times, and incubated for 2 h at 4°C with gentle agitation. The SN was collected using 800 µl centrifuge columns, and the beads were washed 2-times with 400 µl PBS and the flow-through was combined with the SN. The resulting Fab suspension was concentrated on Amicon Ultra-0.5 centrifuge filters with 10 kDa MWCO (Millipore) to 1 µg/µl and validated by SDS-PAGE and coomassie blue staining.

### Antibody labelling

Full length antibodies and Fabs were labelled with Alexa Fluor 488 (A488) or Alexa Fluor 647 (A647) using the Alexa Fluor Antibody Labeling Kit (#A20181; #A20186; Thermo Fisher Scientific) as previously described ^33^, with modifications. 100 µl of 1 µg/µl antibody in PBS and 0.1 M sodium bicarbonate was mixed with the appropriate dyes and incubated at RT for 1 h on a shaker protected from light. Non-conjugated dyes were removed by 3 rounds of buffer exchanges using PBS and Amicon Ultra-0.5 centrifuge filters with 10 kDa MWCO (Millipore). The degree of labelling was typically between 4-7 dyes/antibody molecule which was measured by Nanodrop 2000 (Thermo Fisher Scientific) and calculated according to the formula provided by the Antibody Labeling Kit’s protocol.

### Enzyme linked immunosorbent assay (ELISA)

For the ELISA assays, microtiter wells (ThermoFisher Scientific) were coated with 2 µg/ml of phosphorylated or non-phosphorylated peptide in PBS overnight at 4 °C. The antibodies were diluted in PBS and following incubation for 1h at RT they were revealed with alkaline phosphatase-conjugated goat anti-mouse IgG (Jackson Immunoresearch Laboratories). After several washes with PBS containing 0.05% Tween-20 and addition of pNPP substrate (Sigma-Aldrich), the optical density was measured at 405 nm in an ELISA reader.

### Fluorescent labelling of endogenous proteins by antibody electroporation

Electroporation was performed as previously described ^33^ using the Neon Transfection system with the 10 µl Neon electrode tips (MPK5000 and MPK1096, Thermo Fisher Scientific). Briefly, cells were washed once with Dulbecco’s PBS (DPBS, D8537-500ML; Sigma-Aldrich) and then trypsinized (0.25% trypsin-EDTA; 25200-056; Gibco) for 3 min at 37°C. Detached cells were collected in medium and washed once with DPBS. Cells were then resuspended in electroporation buffer (R) to obtain 10^5^/10 μl cell suspension. 10^5^ cells were mixed with 1 μl of labelled antibody solution (1 µg/µl) and electoporated in an electrode tip using the following settings: 1550 V, 3 pulses, and 10 ms per pulse. The electroporated sample was transferred into 4 ml pre-warmed medium lacking antibiotics in round-bottom tubes (352054; Falcon) and allowed to recover for 30 min in the incubator. Cells were then pelleted, resuspended in 300 μl of pre-warmed medium without antibiotics and put into one well of an 8-well microscopy slide (#80827; Ibidi) for 24 hours at 5 % CO_2_ and 37°C.

### Fluorescence loss in photobleaching (FLIP)

FLIP assays were performed in *U2OS-med* cells expressing the mCherry-tTA-ER activator to measure activator residence times. For the measurements of Pol II, Pol II pSer5, TFIIB and TFIIF residence times, *U2OS-med* cells electroporated with A488-labelled specific antibodies were used. Cells were seeded on Ibidi µ 8-well glass bottom slides and treated with 1 µM 4-OHT overnight. The medium was replaced by phenol red-free medium containing 1 µM 4-OHT before imaging. Microscopy was performed using a Nikon Ti-E inverted microscope with Perfect Focus System equipped with a CSU X1 Yokogawa confocal spinning disc unit, a Leica 100x HC PL APO oil NA 1.40 objective and a Photometrics Prime 95B sCMOS camera. 491 nm and 561 nm lasers were used to excite and photo-bleach the A488 and mCherry flourophores, respectively, and fluorescence was detected using 525/50 and 605/64 band pass filters, respectively. Cells were maintained at 37°C in 5% CO_2_ using a Tokai Hit INUBG2E-TIZ stage top incubator. To perform FLIP the iLAS MODULAR system (Gataca Systems) was used: first, a circular ROI of 125 pixel diameter was defined such that it was as far as possible from the mCherry foci while covering the largest possible area of the same nuclei. Time lapses were obtained by first recording a five frame ‘pre-sequence’ (1.6 s duration / 400 ms interval / 100 ms exposure) followed by 145 cycles of sequential photobleaching (two scans of the ROI with maximum laser power for ∼726 ms followed by a 100 ms exposure image acquisition) and ‘post-sequence’ steps (400 ms duration / 400 ms interval / 100 ms exposure). These produced ∼4.5 min long movies with 295 image frames. For Pol II, Pol II pSer5, TFIIB and TFIIF measurements pre-FLIP images in both the A488 and mCherry channels were also recorded to verify co-localization at the selected foci.

FLIP image sequences analysis and fluorescence were done using a custom ImageJ script. Frames 5-295 were maximum intensity projected and a first ROI enclosing the signal trail of the selected foci (‘spot’) was carefully defined. This ROI was then dilated by 3 pixels to obtain an offset band around the signal to measure the local background (‘locBkgd’). Three additional ROIs of the same size and shape as the first was defined on the same field of view: one in the nucleus of a non-photobleached control cell (‘ctrl’) to correct for time-dependent photobleaching during the course of imaging. The fourth was defined inside the photobleached ROI to confirm photobleaching efficiency and the fifth on a region not covered by cells to measure the experimental background (‘bkgd’). Total fluorescence intensities (‘I’) were measured from every frame from each ROI with areas (‘A’). Relative normalized fluorescence (‘normI’) from the selected foci were computed by first calculating the corrected fluorescence intensities (‘corrI’) of the transcription foci (‘spot’) (1) and the control nuclei (‘ctrl’) (2) from each frame (‘f’). The relative corrected fluorescence of the selected foci was then normalized by photobleaching correction based on the control nuclei (3).

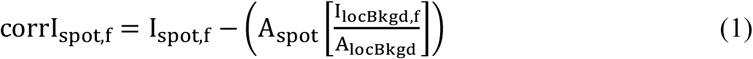

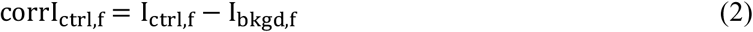

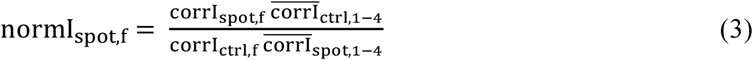

Then both of the following single-(4) and double-component models (5) were fit to each relative normalized fluorescence trace using the curve_fit function of the scipy.optimize package in python.

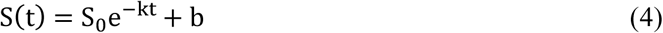

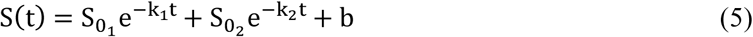

Expected initial values were set as follows: S_0_=1, k=0.02, b=0, 𝑆_01_=1, 𝑆_02_=1, k_1_=0.02, k_2_=0.04, b=0. Bounds were set to 0 to infinite for S_0_, 𝑆_01_, 𝑆_02_, k, k_1_, k_2_. Relative errors of the fits for *k*, *k_1_* and *k_2_* were calculated by dividing the square root of the estimated approximate covariance by the fitted values. Traces with relative errors above 0.33 were excluded from further analyses.

### Confocal microscopy

For live confocal microscopy cells were grown on 8-well microscopy slides in 300 µl medium. The medium was replaced by phenol-red-free free medium supplemented with 2 µg/ml Hoechst33342 prior microscopy. Cells were imaged on the Nikon Ti-E system described above. 405 nm and 562 nm laser lines were used to image the Hoechst33342 nuclear stain and the mCherry at 100 ms exposure times, respectively. 7 z-stacks with 0.5 µm spacing were recorded.

### Time-laps live imaging of induced transcription from the gene array

Cells were imaged on 8-well microscopy slides in 300 µl medium without phenol-red using a Leica DMI8 inverted microscope with Adaptive Focus Control and Z Piezo stage equipped with a CSU-W1 Yokogawa confocal spinning disc scanner unit, a Leica 63x HCX PL APO Lambda blue oil NA 1.40 objective and a Hamamatsu ORCA-Flash4.0 digital CMOS camera. 488 nm and 561 nm lasers and 525/50 and 609/54 band pass filters were used to excite and detect A488 and mCherry, respectively. Cells were maintained at 37°C in 5% CO_2_ with an Okolab bold line H301-mini stage top incubator. Images were acquired in 7 z-stacks with 1 µm spacing with sequential 150 ms exposures in both channels at every z position. 4-5 stage positions were recorded every 30 seconds for 60 minutes streaming fluorescence and z-positions to the camera’s memory. 4-OHT-treatement (1 µM) was performed on stage by pausing image acquisition after recording two frames, and adding 100 µl 4 µM 4-OHT in medium to the cells. Time laps image sequences were processed using a custom tracking and image analyses pipeline.

Time laps image analyses and signal quantification: The ‘x-y’ positions of the transcription sites were tracked in 561 nm channel using the semi-automatic mode of the TrackMate plugin in ImageJ (“Manual tracking with TrackMate”). First, the z-stacks of the raw mCherry image sequences were maximum-intensity projected, 16-fold down-sampled for faster tracking and reversed in time in ImageJ. Then transcription foci were manually identified (mostly one, rarely two per nuclei) and automatically tracked over time in batch (“TrackMate tools”; Quality threshold: 0.001; Distance tolerance: 1; Max nFrames: 121). The resulting tracks were manually curated and adjusted using the “TrackSchemes” tool before the tracked x-y coordinates, the radii (r) of the tracked areas and the corresponding time points (t) were recorded. 3D tracking (i.e., identification of the ‘z’ positions of each spot at each tracked time points) was done using a custom ImageJ script as following: a circle ROI with x-y coordinates and r radius for every t was mapped back on the raw image stacks in the 561 nm channel and the z-frame with the highest total pixel intensity under the ROI was selected with the constrain that the z-position cannot move more than 1 frame between time points. The first detected (i.e. earliest) x-y-z coordinates were propagated to the frames preceding the time frame of the first detection to have consistent movie lengths and record fluorescent data for the entire time courses. To trace the outlines of the spots, 40×40 pixel regions centered on the x-y-z-t coordinates were cropped from the raw 561 nm channel movies for every track and binary masks covering the precise shape of the spots were calculated for every time point using a convolutional neural network-based machine learning algorithm in R. Masks were further processed in ImageJ using a custom script: first to avoid wrongly identified masks, consecutive masks between t and t+1 timeframes were required to overlap with at least one pixel, otherwise the t mask was replaced by the t+1 mask. The remaining correct masks were then converted to ‘signal’ (S) ROIs which were mapped back on the original raw image stacks (x-y-z-t), then enlarged by two pixels and used to collect total fluorescent intensities from the transcription site. Each of these signal ROIs were dilated by 7 pixels to obtain an offset band surrounding the signal ROIs and were set as ‘local background’ (locBkgd) ROI to collect local background fluorescence around the transcription site. Total fluorescence intensities (I) and areas in pixels (A) were measured for each ROI over time. A 50×50 pixel rectangle image stacks around the center of each signal ROIs were also cropped from both channels for visual curation of the transcription sites. A photobleaching correction factor (F_t_) was calculated frame-by-frame for both channels by dividing total pixel intensities of maximum intensity-projected field of views from consecutive time points (F_t_=totI_t-1_/totI_t_). Normalized total intensity for each transcription site was calculated over time by subtracting background fluorescence from the total signal and corrected for photobleaching:

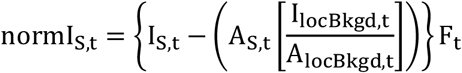

The first activator binding events (t_0_) were determined as the time points of the first significant increase of mCherry signal at the transcription site over the local background. A custom python script was used to perform Mann-Whitney U (MWU) test (scipy.stats.mannwhitneyu) for each time point with a 10 frame forward moving window comparing signal and local background mean raw fluorescent intensities. Then t_0_ was set as the first t_n_ time point for which: 1) MWU p<0.01; 2) for at least 7 of the next 9 time points (t_n+1_-t_n+9_) p<0.01 as well (i.e. the signal remains significantly higher than background with little fluctuation); 3) the five frame forward moving average signal at t_n_ (t_n_-t_n+4_) was above background; 4) the five frame forward moving average signal for t_n_ was at least double of that of t_n-5_ (i.e. the signal started to increase after t_n_); 5) the forward moving background-corrected average signal for t_n+5_-t_n+19_ was at least 40 % higher than for t_n_-t_n+4_ (i.e. the signal kept an increasing trend over 20 frames). The same approach was also used to determine the first significant t_0_ detection of the antibody-labelled endogenous factors (Supplementary Fig. 3g, h) Individual raw mean signal and background traces were visually curated, and wrong t_0_ detections were manually corrected and low-quality tracks were excluded from further analyses.

### Immunofluorescence (IF) analyses

Cells were seeded in 12-well plates onto round glass coverslips. Next day after removing the medium the cells washed twice with PBS, then fixed for 5 min in 4% PFA (Electron Microscopy Sciences) in PBS pre-warmed to 37°C, and washed again twice with PBS. Permeabilization was performed for 20 min at RT by in 0.1% Triton X-100 in, followed by two washes with PBS. The cells were then incubated with 2 µg/ml primary antibody in 1% BSA in PBS for 1 h at RT, then washed two times for 5 min with 0.02% Triton X-100 in PBS and once with PBS. Alexa Fluor 488–conjugated goat anti-mouse secondary antibody (Thermo Fisher) was added to the cells at 1/3000 dilution in 1% BSA in PBS for 1 h, then the cells were washed two times for 5 min with 0.02% Triton X-100 in PBS and once with PBS, and mounted using Vectashield containing DAPI. Stained cells were imaged on a Leica DMI8 spinning disc confocal microscope as described above, using a Leica 100x HCX PL APO oil NA 1.47 objective, and 405 nm and 488 nm lasers for DAPI and A488 excitation, respectively, and images with eleven z-stacks with 0.2 µm spacing were recorded. Custom Fiji/ImageJ script was used for the quantification of nuclear fluorescence from the A488 channel on maximum intensity z-projected images after background subtraction and nuclear segmentation (based on the DAPI signal).

### Direst stochastic optical reconstruction microscopy (dSTORM)

U2OS-med cells were seeded on Ibidi µ 8-well glass bottom slides and treated with 1 µM 4-OHT overnight. Cells were washed twice with PBS then fixed with 200 µl 37°C pre-warmed 4% paraformaldehyde (#15710 Electron Microscopy Sciences) in PBS for 5 min at RT and washed again twice with PBS. Permeabilization was performed with 200 µl 0.1 % Triton X-100 in PBS for 20 min at RT followed by two washes with PBS. Cells were then stained with 2 µg/ml A647-labelled anti-ER or anti-Pol II antibodies for 1 hour at RT in 200 µl 1 % BSA in PBS, and washed twice with 0.02 % Triton X-100 in PBS and once with PBS. dSTORM was performed as previously described ^69^. Samples were mounted directly before imaging by filling up the well with ∼800 µl imaging buffer (200 mM Tris-HCl pH 8.0, 10 mM NaCl, 10% w/v glucose, 200 U/ml glucose oxidase [G2133, Sigma-Aldrich], 1000 U/ml catalase [C1345, Sigma-Aldrich], 50 mM mercaptoethylamine [30080, Sigma-Aldrich], 2 mM cyclooctateraene [138924, Sigma-Aldrich]) and covering the top with a clean coverslip air-tight, avoiding air-bubbles. Imaging was performed on a modified Leica SR GSD system equipped with an HCX PL APO 160x/1.43 Oil CORR TIRF PIFOC objective (Leica), a custom double-band filter cube and an Andor iXon Ultra 897 (DU-897U-CS0-#BV) EMCCD camera. For fluorophore excitation a 642 nm 500 mW fiber laser (MBP Communication Inc.) and a 405 nm 50 mW diode laser (Coherent Inc.) was used ^69^.

Samples were first inspected in normal fluorescence mode and cells with clear nuclear mCherry foci were selected and reference images with 532 nm and 674 nm laser excitation were acquired in both channels. Then mCherry was photobleached with high intensity laser power to avoid signal bleed through. Samples were then imaged in super resolution mode: the sample was illuminated by 40% laser power using the 642 nm laser at 10 ms exposure time. Acquisition was started manually after the appearance of separate single-fluorophore events. When the number of localizations started to drop (10-20 min), additional illumination with the 405 nm laser using gradually increasing intensity was applied for fluorophore reactivation. Typically, 70-100 thousand images were acquired until near complete photobleaching of the fluorophore. Localization tables from the Leica Las X software were exported and further processed for drift corrections and localization filtering using 400 photons threshold with the SharpVisu Matlab workflow ^70^. The resulting 7-10 nm pixel size super-resolution images are 2D histograms of the single-molecule coordinates.

## Acknowledgements

We thank Susan M. Janicki for kindly providing the U2OS 2-6-3 cell line. We thank also Elvire Guiot and Erwan Grandgirard at the Imaging Centre of IGBMC, Rafael Schoch at the Super Resolution Microscopy platform of IGBMC, and Claudine Ebel and Muriel Philipps at the CytoEast facility for their expert technical assistance.

## Author contributions

A.O. N.M. and L.T. conceived and designed the research. A.O., S.C. conducted experiments. M.A. raised and tested antibodies. A.O. O.T. N.M. and L.T. analysed and interpreted the results. N.M. and L.T. supervised the study. A.O. and L.T. wrote the first draft, and N.M. A.O. and L.T. finalized the manuscript.

## Supplementary Figures

**Supplementary Fig. 1.**
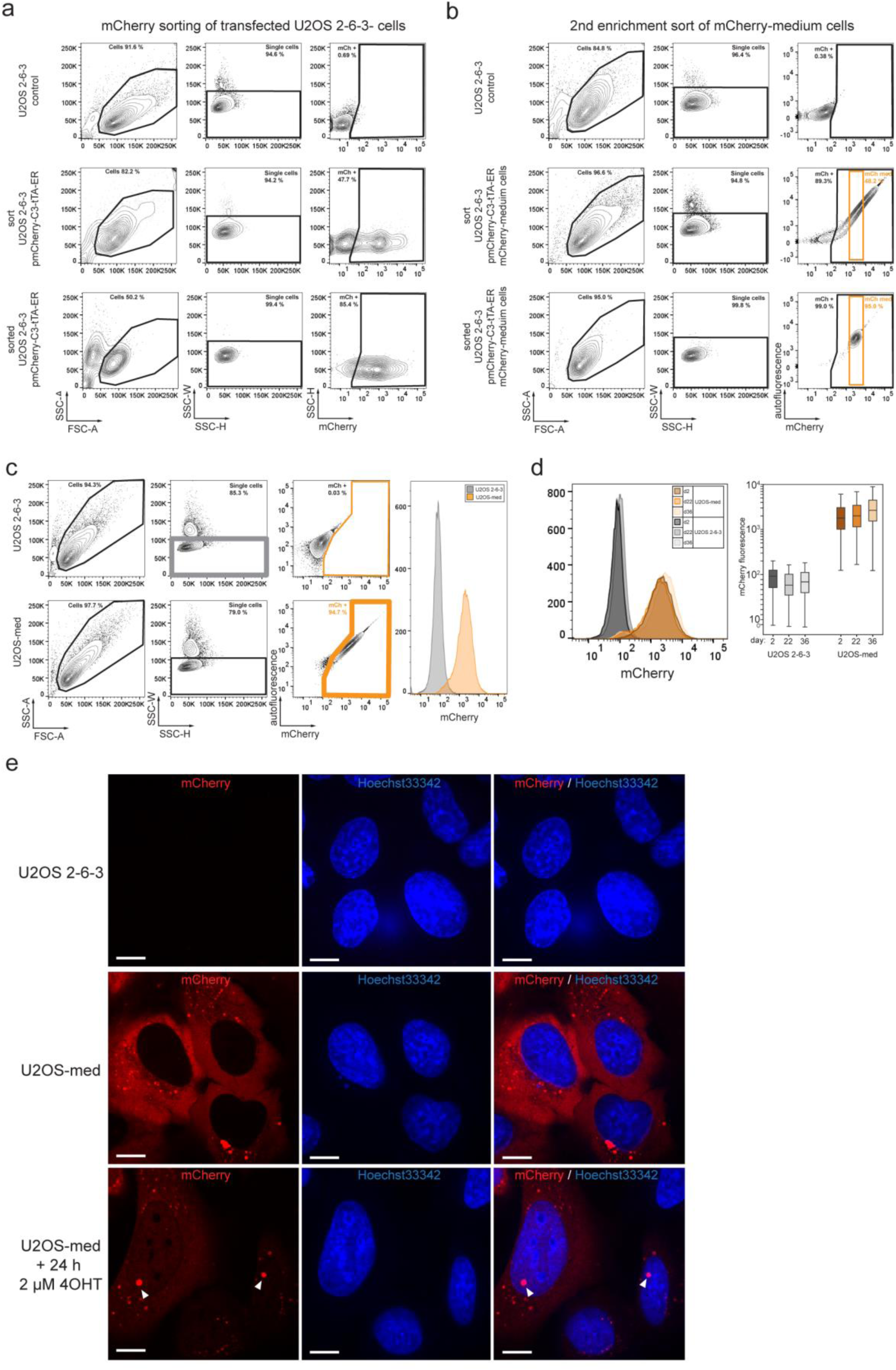
Generation and characterization of the U2OS-med cell line. **a** U2OS 2-6-3 cells were electroporated with the pmCherry-C3-tTA-ER vector, and mCherry positive cells were sorted 24 h later on FACSAria II. Gating strategy and the corresponding percentage of sub-populations are shown: mCherry expression (right) was analysed on the single cell subpopulation (middle) gated from the total intact cells (left). Top row shows non-transfected negative control cells. Middle row shows transfected cells which were sorted on the mCherry-positive (mCh +) gate. Bottom row shows the analyses of the sorted cells. The mCherry positive sorting gate was defined to exclude most negative control single cells (top right). **b** Top row shows U2OS 2-6-3 control cells. Middle row shows the second re-sorting of the expanded cells derived from (**a** bottom). The sorting gate (mCh med; orange) is set to isolate single cells with medium levels of mCherry expression. Other gates are set as in (**a**). Bottom raw shows the analyses of the sorted cells. **c** Flow cytometry analyses of the established U2OS-med cell line using LSRFortessa. Gating strategy as in (**a**). The histogram on the right shows the distribution of mCherry fluorescence intensities from control single cells (thick gray gate) and U2OS-med single cells expressing mCherry (thick orange gate). **d** Stable mCherry-tTA-ER expression in U2OS-med cells. U2OS-med cells were grown for 36 days in culture over 10 passages. Samples were analysed by flow cytometry using LSRFortessa at days 2, 22 and 36. The histogram on the left and the box plot on the right show the distributions of mCherry fluorescence intensities from control U2OS 2-6-3 and U2OS-med single cells gated as in (**c**). **e** U2OS-med cells were treated overnight with 2 µM 4-OHT (bottom) or mock (EtOH; middle) and then imaged live on a Nikon spinning disk confocal microscope. U2OS 2-6-3 control cell are shown on top. Arrowheads point to enriched mCherry accumulation at the gene arrays on the background of increased nuclear mCherry signal after 4-OHT treatment. Scale bars are 10 µm. SSC: side-scatter; FSC: forward-scatted; A: area; W: width; H: height.

**Supplementary Fig. 2.**
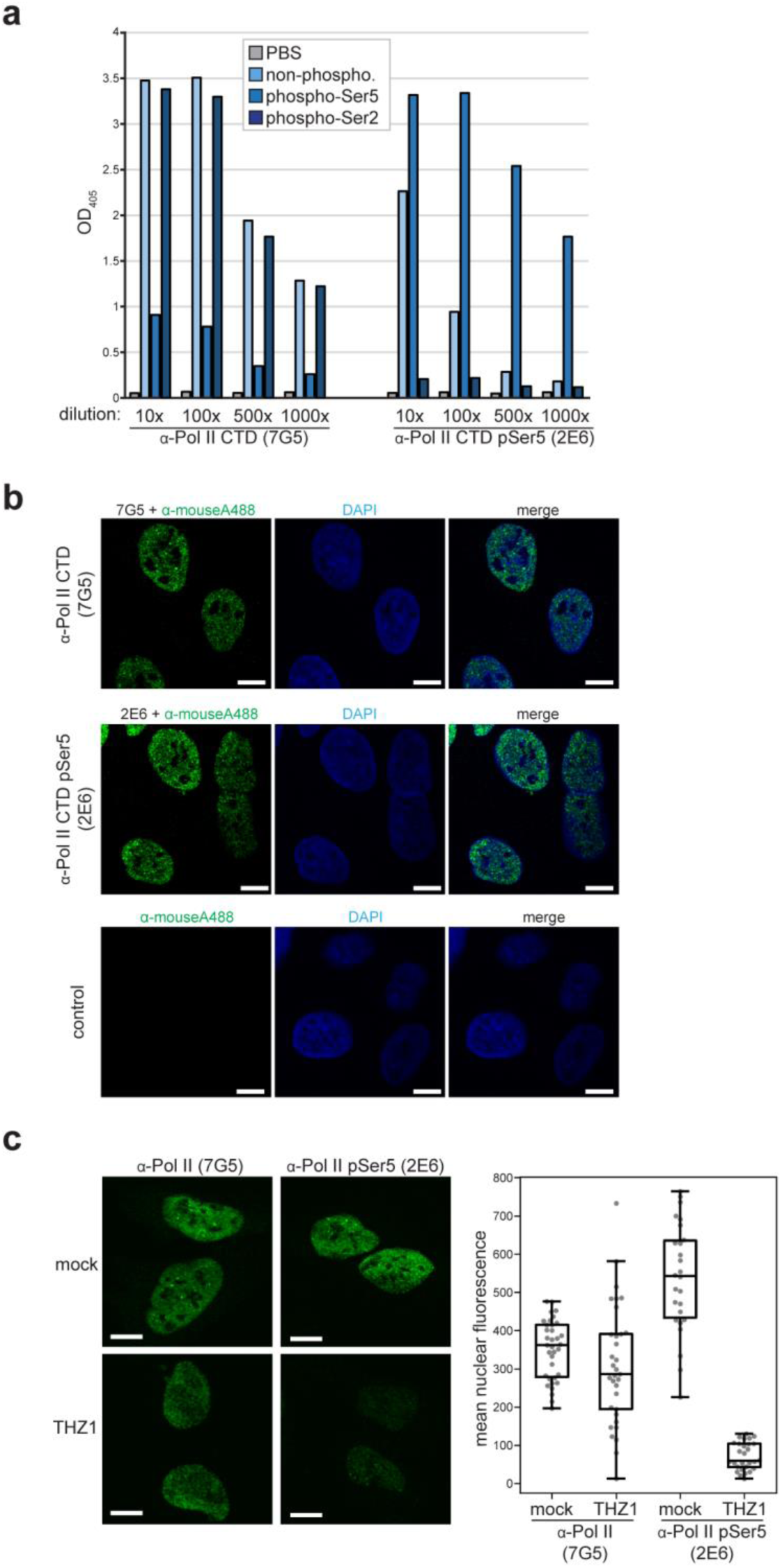
Characterization of the newly generated anti-Pol II CTD pSer5 antibody. **a** Peptide ELISA assays showing the Ser5 phosphorylated heptapeptide repeat-specificity of our newly generated anti-Pol II CTD pSer5 mouse monoclonal antibody (mAb 2E6). Sequential dilutions of culture supernatants from hybridoma cells producing either anti-Pol II CTD (clone 7G5) or our newly generated anti-Pol II CTD pSer5 (clone 2E6) antibodies were tested on ELISA plates coated with either PBS, or non-phosphorylated peptides, or peptides phosphorylated at Ser5 or at Ser2 residues, as indicated. **b** Immunofluorescence showing similar Pol II staining patterns with the anti-Pol II CTD (7G5) and the anti-Pol II CTD pSer5 (2E6) antibodies: predominantly nuclear staining with enriched fluorescence at sub-nuclear foci. Fixed and permeabilised U2OS 2-6-3 cells were stained with purified anti-Pol II (top) or anti-Pol II pSer5 (middle) primary, and AlexaFluor488-labelled anti-mouse secondary antibodies, and subsequently imaged on a Leica spinning disc confocal microscope at 100x magnification. DAPI was used for nuclear staining. The bottom row shows negative control staining with the secondary antibody alone. Images are representative of four experiments. Single z-planes are shown. Scale bars are 10 µm. **c** Inhibition of Pol II CTD-phosphorylation at Ser5 with THZ1 strongly decreases IF staining with the anti-Pol II CTD pSer5 antibody. U2OS cells were treated with either the Pol II CTD pSer5-specific inhibitor THZ1 (15 µM) or by DMSO (mock) for 2 hours prior fixation, permeabilisation, staining with either anti-Pol II CTD (7G5) or anti-Pol II pSer5 (2E6) mAbs and imaging as in (**b**). Images of maximum intensity-projections of seven 0.2 µm z-stacks are shown on the left as indicated. Distributions of the mean nuclear fluorescence intensities measured after nuclear segmentation and background subtraction is show on the left. Results are representative of two experiments. Scale bars are 10 µm.

**Supplementary Fig. 3.**
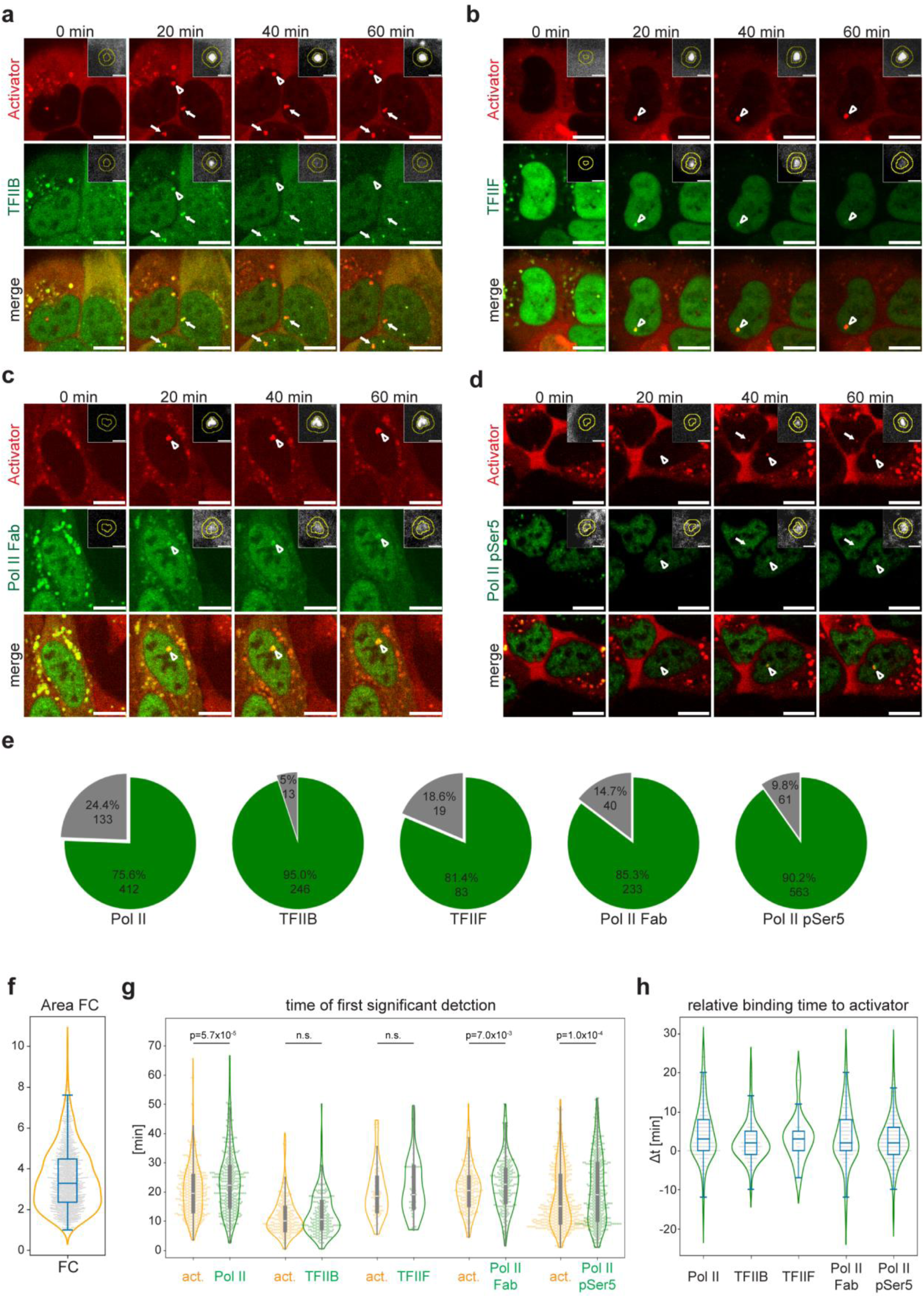
Characterization of activator, Poll II and GTF recruitment dynamics. **a**-**d** Representative snapshot images at the indicated time points from of U2OS-med cells that were labelled with anti-TFIIB (**a**), anti-TFIIF (**b**), anti-Pol II Fab (**c**) or anti-Pol II pSer5 (**d**) antibodies conjugated with Alexa488 overnight, activated with 4-OHT and imaged for 1 h. Arrows point to the co-localisation (merge; bottom row) of the activator (red; top row) and the indicated labelled endogenous protein (green; middle row) signals at the activated transcription sites. Arrow heads show the ones that were tracked as shown in the insets. Scale bars are 10 µm. Insets show the tracking of the signal (inner circle) and local background (outer ring) areas highlighted by the arrowheads. Inset scale bars are 2 µm. **e** Numbers and proportions of the detected induced activator binding which had (green; n=1537) or did not have (grey) overlapping signal from A488-labelled recruitment of anti-Pol II, anti-TFIIB, anti-TFIIF, anti-Pol II Fab or anti-Pol II pSer5 at the transcription site. **f** Fold change distribution of maximum activator signal area over the activator signal area at the time of first detected activator binding (t_0_) at the 1537 transcription sites measured in Fig. 1f. Median FC = 3.286 (95% CI 3.196-3.385). **g** Pairwise comparisons of the time points (min) of first significant detection of the activator (orange) and the indicated endogenous factors (green) at the activated transcription site. Pairwise median and (95% CI) values in minutes: activator / Pol II=19.5 (18.5-21.0) / 22.25 (21.0-23.75); activator / TFIIB=10.0 (9.0-11.0) / 10.5 (9.5-12.0); activator / TFIIF=18.5 (15.5-20.0) / 19.0 (16.5-23.25); activator/Pol II Fab=20.5 (19.5-21.5) / 22.5 (21.5-24.0); Pol II pSer5= 15.0 (14.0-16.0) / 19.0 (17.0-20.75). p-values correspond to Kruskal-Wallis tests. **h** distribution of times (min) between the first significant detection of the activator signal and that of the indicated factors (Δt=t_0[factor]_-t_0[activator]_) from each trace. Median and (95% CI) values in minutes: Pol II=2.0 (1.5-2.5); TFIIB=1.0 (0.5-1.0); TFIIF=1.75 (1.0-2.5); Pol II Fab=1.0 (1.0-2.0); Pol II pSer5=1.0 (1.0-1.5).

**Supplementary Fig. 4.**
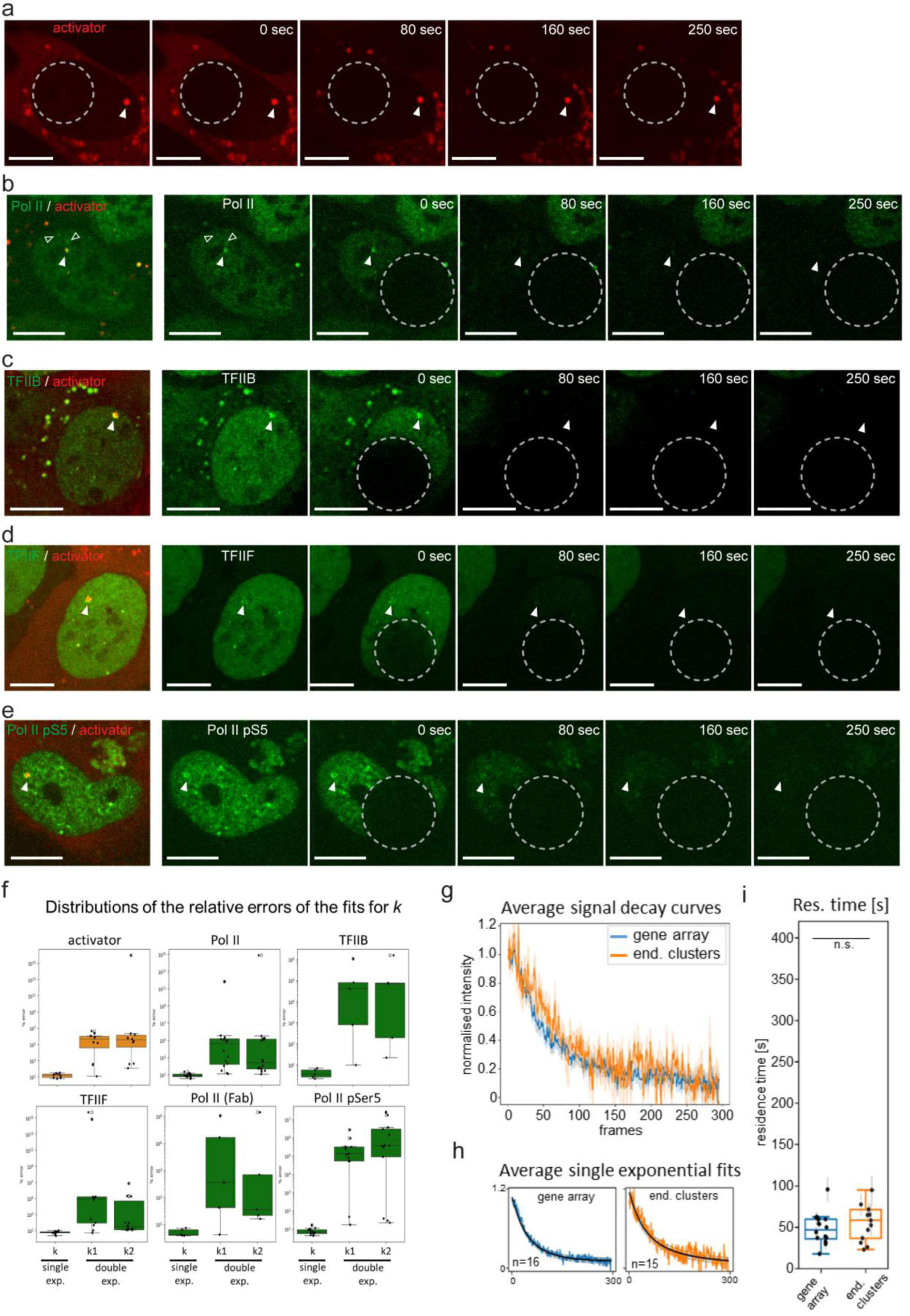
Measurements corresponding to activator, Pol II and GTF FLIP assays. a-e. Example images of the FLIP assay for activator (**a**), Pol II (**b**), TFIIB (**c**), TFIIF (**d**) and Pol II pSer5 (**e**). Filled arrowheads indicate the co-localization of the labelled endogenous proteins and/or activator signals at the transcription sites. Open arrowheads point to examples of endogenous Pol II clusters. **f** Box plots comparing the distributions of the relative errors of the fits for single (k) or double (k1, k2) exponential functions fitted to the FLIP decay curves of the indicated factors from Fig. 3c. **g** Comparison of the average FLIP decay curves (±SEM) of Pol II at endogenous clusters vs the gene array. **h** Average FLIP decay curves from (**g**) and the corresponding average fitted single exponential functions (black curves; ±SEM). **i** Distribution of the residence times (1/k) derived from the fitted dissociation rate constants in (**h**). median±SEmedian [sec] 1/k for Pol II at the gene array (same as Fig. 3c-e): 46.8±5.8; or at endogenous clusters: 55.1±7.1; Error bars: relative errors of the model’s fit for each decay curve. Student’s T-test (p= 0.3369).

**Supplementary Fig. 5.**
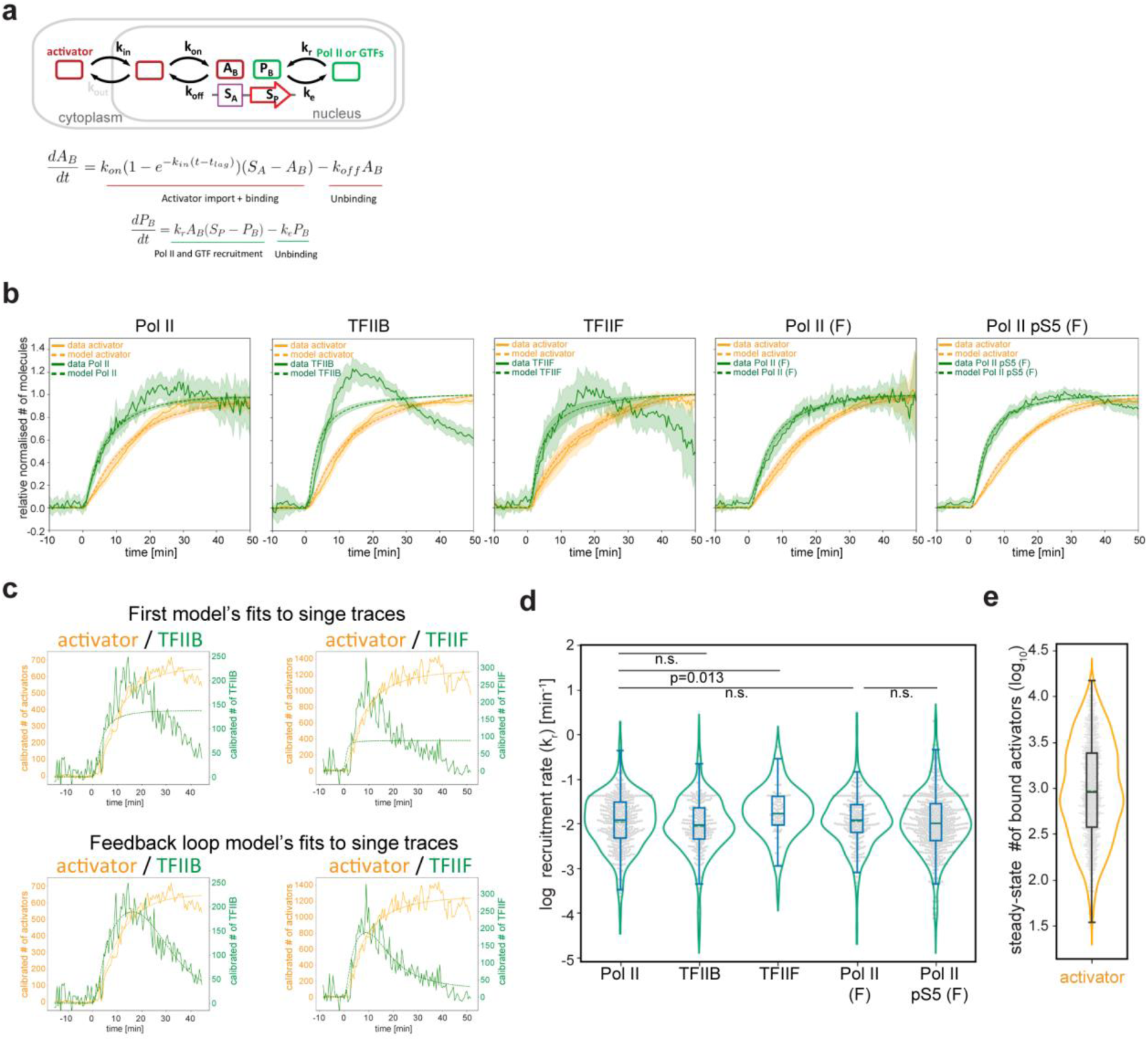
Kinetic models of activator, Pol II and GTF recruitment following transcription activation. **a** Model for the activation-induced recruitment of activator, Pol II and GTFs. **b** Calibrated binding curves and model fits using the model in (**a**) for Pol II, TFIIB, TFIIF, Pol II Fab, Pol II pSer5 Fab (left to right, green) and activator (orange). Continuous and dashed lines show average normalized calibrated data *(±3xSEM)* and the average fitted models *(±3xSEM)*, respectively **c** Single traces of TFIIB (left, green) or TFIIF (right, green) and activator (orange) calibrated binding curves (solid lines) and the corresponding model fits (dashed lines) from the simple model in (**a**) (top) and the updated model in Fig. 4a (bottom). **d** Distributions of model-predicted recruitment rates of the indicated factors after fitting the model in Fig. 4a. median±SEmedian [sec] values for Pol II: 0.0119±0.0041; TFIIB: 0.0091±0.0064; TFIIF: 0.017±0.0099; Pol II Fab: 0.0118±0.0075; Pol II pSer5 Fab: 0.0101±0.0086. p-values were calculated with Holm-Šidák-correction from a post-hoc Dunn test after a significant Kruskal-Wallis test (F=18.11, p=0.001). **e** distribution of the model-predicted steady-state number of bound activators: (k_on_S_A_/[k_on_+k_off_]). n=1537; median=901.21 [95% CI 827.95-993.55].

**Supplementary Fig. 6.**
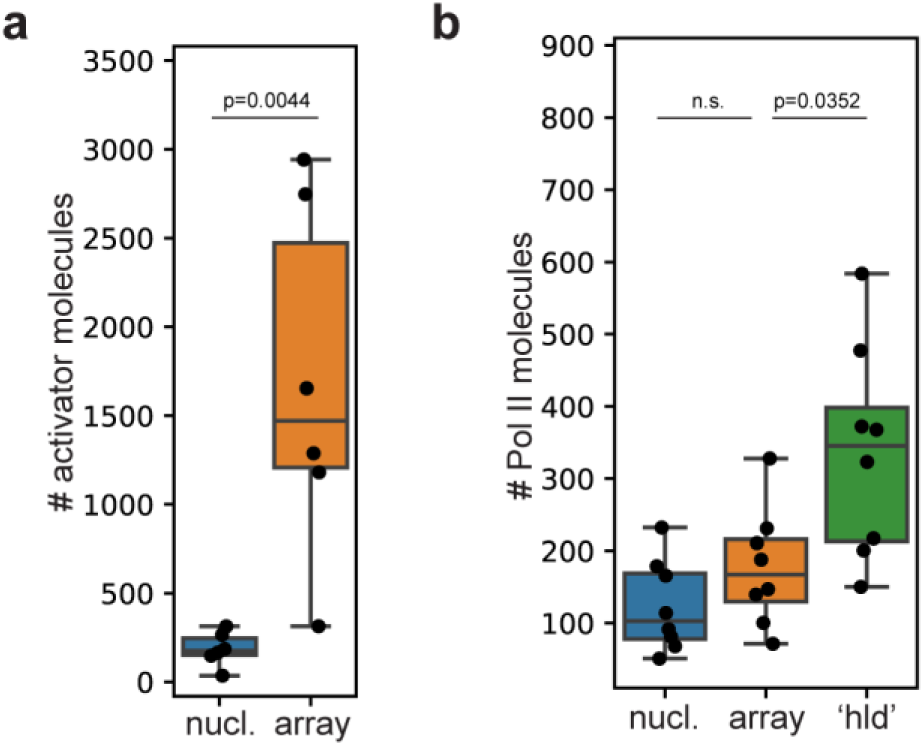
Super resolution microscopy of the activator and Pol II. a, **b** Quantification of dSTORM images as in Fig. 5. Box plots showing the distribution of the calculated total number of activator (**a**) (n=6) or Pol II (**b**) (n=8) molecules at nucleoplasmic control regions (nucl.), at the gene array (array) or at high local density (hld) regions. Mean±SEM for (**a**) activator array: 1687.52±408.2; activator nucl.: 185.36±39.68; *p*-value is calculated from a Student’s *t*-test. For (**b**): Pol II array: 176.92±28.77; Pol II hld.: 336.57±51.91; Pol II nucl.: 122.68±22.34. p-values were calculated from Student’s t-tests with Bonferroni correction.

**Supplementary Fig. 7.**
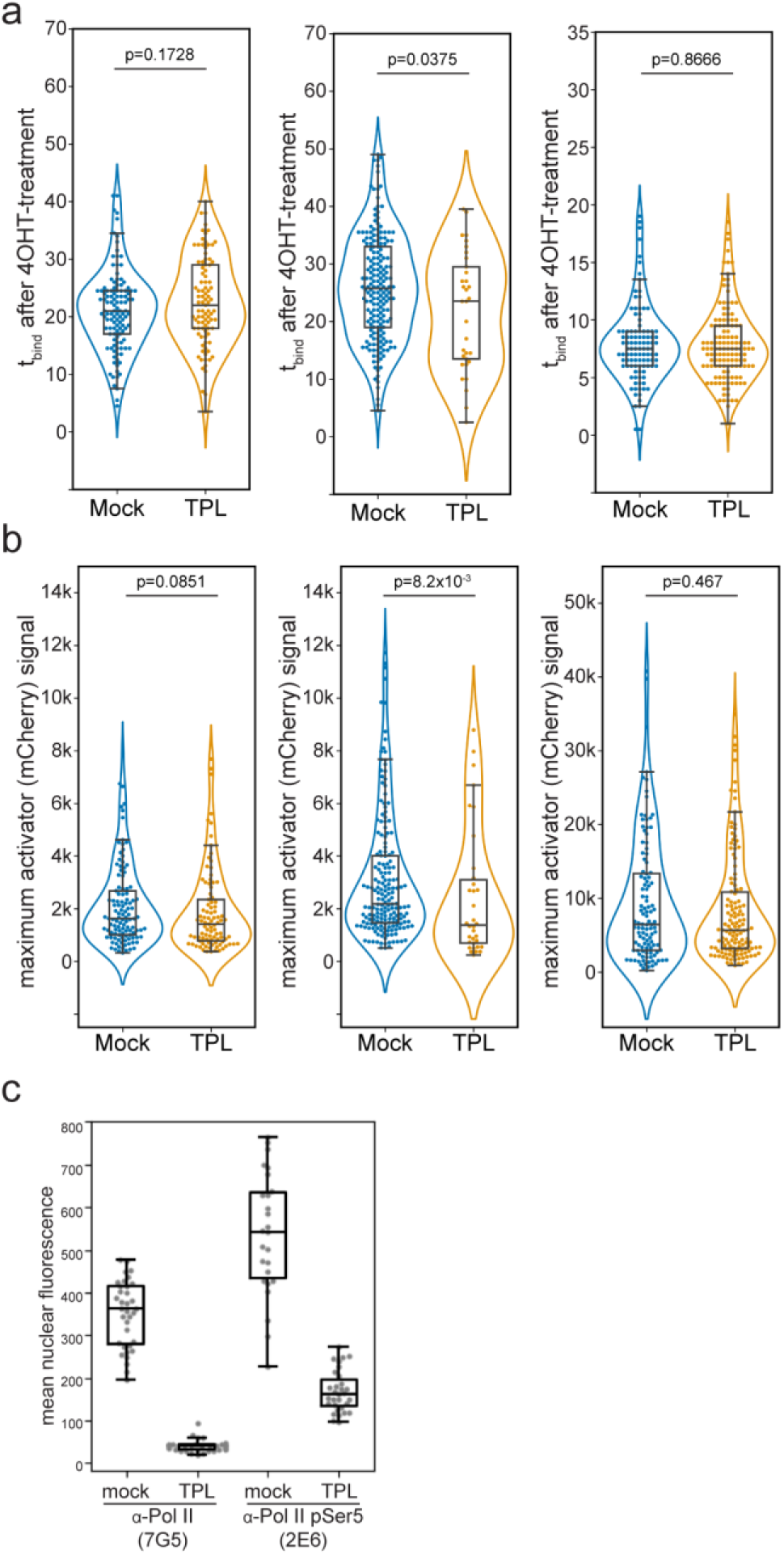
Triptolide (TPL) has no major effect on activator binding. Distributions of **a** activator binding times and **b** maximum activator (mCherry) signal in response to transcription inhibition with TPL from the three different sets of antibody-labeled samples (corresponds to Fig 6a, b). p-values were calculated from Mann-Whitney U tests. **c** 2 h TPL treatment strongly decreases Pol II and Pol II pSer5 levels. U2OS cells were treated with either TPL (5 µM) or by DMSO (mock) for 2 hours and IF was performed using either anti-Pol II or anti-Pol II pSer5 antibodies as in Supplementary Fig. 2b. Boxplots show the distributions of the mean nuclear fluorescence intensities measured as in Supplementary Fig. 2c.

## Supplementary Movies

**Supplementary Movie. 1.** Example time laps recording of live U2OS-med cells electroporated with AlexaFluor488-labelled anti-Pol II antibodies and imaged on a spinning disc confocal microscope the next day for 1 hour with 2/min frame rate. Cells were treated with 4OHT at the 1 min (second frame) time points of image acquisitions. Right segment shows the overlay of the mCherry signal from the activator (left segment) and the A488 signal from the antibody-labelled endogenous Pol II molecules (middle segment). Scale bars are 10 µm. Insets show the tracking of the signal (inner circle) and local background (outer ring) areas at the transcription site. Inset scale bars are 2 µm. Corresponds to Fig. 1b.

**Supplementary Movie. 2** Same as Supplementary Movie 1, except that AlexaFluor488-labelled anti-TFIIB antibodies were used to label endogenous TFIIB molecules. Corresponds to Supplementary Fig. 3a.

**Supplementary Movie. 3** Same as Supplementary Movie 1, except that AlexaFluor488-labelled anti-TFIIF antibodies were used to label endogenous TFIIF molecules. Corresponds to Supplementary Fig. 3b.

**Supplementary Movie. 4** Same as Supplementary Movie 1, except that AlexaFluor488-labelled anti-Pol II Fab antibodies were used to label endogenous Pol II molecules. Corresponds to Supplementary Fig. 3c.

**Supplementary Movie. 5** Same as Supplementary Movie 1, except that AlexaFluor488-labelled anti-Pol II pSer5 fabs were used to label endogenous Pol II molecules phosphorylated on the CTD Serine 5 residues. Corresponds to Supplementary Fig. 3d.

